# Genetic “General Intelligence,” Objectively Determined and Measured

**DOI:** 10.1101/766600

**Authors:** Javier de la Fuente, Gail Davies, Andrew D. Grotzinger, Elliot M. Tucker-Drob, Ian J. Deary

## Abstract

It has been known for 115 years that, in humans, diverse cognitive traits are positively intercorrelated; this forms the basis for the general factor of intelligence (*g*). We directly test for a genetic basis for *g* using data from seven different cognitive tests (N = 11,263 to N = 331,679) and genome-wide autosomal single nucleotide polymorphisms. A genetic *g* factor accounts for 58.4% (SE = 4.8%) of the genetic variance in the cognitive traits, with trait-specific genetic factors accounting for the remaining 41.6%. We distill genetic loci broadly relevant for many cognitive traits (*g*) from loci associated with only individual cognitive traits. These results elucidate the etiological basis for a long-known yet poorly-understood phenomenon, revealing a fundamental dimension of genetic sharing across diverse cognitive traits.

---

Scores on psychometric tests of cognitive abilities (intelligence) robustly predict educational performance, socio-economic attainments, everyday functioning, health, and longevity (*1–3*). In 1904, Charles Spearman identified a positive manifold of intercorrelations among school test results and estimates of intelligence, leading him to propose that they arise from a single general dimension of variation, which he termed general intelligence (and which he denoted as *g*) (*4*). He theorized that most of the remaining variance in each cognitive test was accounted for by a factor specific to that test, which he called *s*. Thus, some variance in each cognitive test is shared with all other cognitive tests (*g*), and some is specific to that test (its *s*). Hundreds of studies have since replicated the finding that, when several or many diverse cognitive tests are administered to a sizeable sample of people, a *g* factor is found that accounts for about 40% of the total test variance (*5,6*). Considerable efforts over the past century have been placed on identifying the biological bases of *g*, spanning levels of analysis from molecular, to neuroanatomical, to cognitive (*7–11*).

Psychometrically, a hierarchical structure of cognitive abilities is commonly agreed, with cognitive tests’ variance accounted for by three different strata of variation (Fig. S1), representing each test’s specific variance (*s*), broad domains of cognitive function (e.g. reasoning, processing speed, memory), and *g* (*5*). Most twin studies that examine the heritability of human intelligence differences use test scores that mix *g* and *s*; relatively few have applied the twin method to the hierarchy of cognitive test score variance (*12*). The twin-based studies that have separated *g* variance from *s* variance indicate a strong heritable basis for *g*, suggesting that cognitive traits are positively correlated substantially because of strongly overlapping genetic architecture (*13–17*). These findings are consistent across studies, but draw inferences only indirectly, via comparisons of phenotypic correlations across monozygotic and dizygotic twins. To date, molecular genetic methods (GWAS) have been applied to phenotypic indices of *g* (*18–20*). However, the cognitive measures used have been scores derived from individual tests, principal components analysis of several tests, or composite scores from omnibus cognitive tests. They therefore have mixed *g* and *s* variance. On the other hand, GWASs of individual cognitive tests have not taken into account that all tests contain *g* variance, and so genetic loci for, say, verbal declarative memory or processing speed might either be related to *g* and/or to the named cognitive property (*21,22*). This is a common error in both phenotypic and genetic cognitive studies (*23*). Here we sought to test for a genetic *g* factor directly, using a multivariate molecular genetics approach that is applied to the hierarchy of cognitive variation. This investigation provides key insights into the shared genetic architecture across multiple cognitive traits and allows the explicit identification of genetic variants underlying *g*. It distinguishes variants broadly relevant for many cognitive traits (via genetic *g*) from variants associated with only individual cognitive traits (via genetic *s* factors).

## Results

Data for the present study came from the UK Biobank, a biomedical cohort study that collects a wide range of genetic and health-related measures from a population-based sample of community-dwelling participants from the UK. Participants were measured on up to seven cognitive traits using tests that show substantial concurrent validity with established psychometric tests of cognitive abilities, and good test-retest reliability (*24*): *Reaction Time* (RT; *n* = 330,024; perceptual motor speed)*, Matrix Pattern Recognition* (*n* = 11,356; nonverbal reasoning), *Verbal Numerical Reasoning* (VNR; *n* = 171,304; verbal and numeric problem solving; the test is called ‘Fluid intelligence’ in UK Biobank), *Symbol Digit Substitution* (*n* = 87,741; information processing speed), *Pairs Matching Test* (*n* = 331,679; episodic memory), *Tower Rearranging* (*n* = 11,263; executive functioning), and *Trail Making Test* – *B* (Trails-B; *n* = 78,547; executive functioning). A positive manifold of phenotypic correlations was observed across the seven cognitive traits (Table S2).

We first aimed to determine the genetic contribution of *g* to the variation in each of the cognitive traits using molecular genetic data. We used a multivariable version of Linkage Disequilibrium Score Regression (LDSC) (*25*) implemented in Genomic SEM (*26*) to estimate genetic correlations among the cognitive traits from molecular genetic data. Prior to this formal modelling, we conducted exploratory analyses on the cognitive traits’ genetic correlations, similar to those often conducted on cognitive phenotypes.

As was first reported at the phenotypic level by Spearman 115 years ago (*4*), we identified a positive manifold of *genetic* correlations among the UK Biobank cognitive traits, ranging from .14 to .87 (*M* = .53, *SD* = .22; Fig. S2; Tables S1–S2). The mean genetic correlation was .530, and the first principal component accounted for a total of 62.17% of the genetic variance. Using genomic-relatedness based restricted maximum-likelihood (GCTA-GREML) (*27,28*), a different estimator of the genetic correlations among the seven cognitive traits (Fig. S3), the mean genetic correlation was .502, and the first principal component accounted for 61.24% of the genetic variance. The correlation between LDSC- and GCTA-GREML-derived genetic correlations was r = .946, indicating very close correspondence between results of the two methods (Fig. S4).

We then proceeded with Genomic SEM to model the genetic covariance matrix formally; this allowed us to evaluate the fit of the genetic *g* factor model, estimate SEs for model parameters, estimate genetic correlations with collateral phenotypes, and incorporate genetic *g* explicitly into multivariate discovery. We applied Genomic SEM to fit a single common factor model to the LDSC-derived genetic covariance matrix among the seven cognitive traits. This model specified the genetic component of each cognitive trait to load on a single common factor, which we term genetic *g*. For each trait, we additionally estimated residual, trait-specific genetic variance components (genetic *s*s). Thus, we formally distill the molecular genetic contributions of *g* and *s* to heritable variation in each of the cognitive traits, and test the fit of this model. Fig. 1 displays the standardized estimates for this model (top panel) and the standardized estimates from a phenotypic factor model (bottom panel) fitted to the phenotypic covariance matrix (Fig. S5; Table S3). Standardized genetic factor loadings were linearly associated with standardized phenotypic factor loadings and generally higher in magnitude (Fig. S6). Table 1 additionally reports both the proportion of genetic *g*:genetic *s* variance for each cognitive trait, and the respective absolute contributions. The genetic *g* factor accounted for 58.37% (SE = 4.84%) of the genetic variance in the seven cognitive traits. All of the traits’ loadings were statistically significant, ranging from. 31 to .98 (*M* = .74, *SD* = 0.22). Four of the cognitive traits have a genetic contribution to their variance that is principally from genetic *g* and much less from a genetic *s*; these are Trails-B (95.30% genetic *g*; 4.70% genetic *s*), Tower (72.80% genetic *g*; 27.20% genetic *s*), Symbol Digit (69.10% genetic *g*; 30.90% genetic *s*), and Matrices (68.20% genetic *g*; 31.80% genetic *s*). VNR (51.40% genetic *g*; 48.60% genetic *s*) and Memory (42.40% genetic *g*; 57.60% genetic *s*) are more evenly split. RT has the majority of its genetic influence from a genetic *s* (9.50% genetic *g*; 90.50% genetic *s*). We emphasize one important implication of these results, i.e. that genetic analyses of some of these individual traits will largely reveal results relevant to *g* rather than to the specific abilities thought to be required to perform the test (*23*). Fit indices (χ^2^(14) = 117.019, *p* < .0001; CFI = .970; SRMR = 0.088) indicated that the factor model closely approximated the observed genetic covariance matrix (Figs. S7–S8). As the pre-specified model was parsimonious and the fit was close, we chose to forego implementing data-driven exploratory steps to further improve fit. Tables S4–S5 report full parameter estimates for genetic and phenotypic factor models.

**Table 1.** Common factor solutions for the genetic (top section) and phenotypic (bottom section) covariance structure of seven UKB cognitive traits.

| Standardized Factor Loadings |  |  | Common (g) and specific (s) sources of genetic variation |  |  |  |  |
| --- | --- | --- | --- | --- | --- | --- | --- |
| Genetic g |  |  | Proportion of genetic variation explained by genetic g and genetic s |  | Proportion of phenotypic variation explained by genetic g and genetic s (HapMap3 Common Variants Only) |  |  |
| Cognitive Trait | Estimate | SE | Common (g) | Specific (s) | Common (g) | Specific (s) | Total SNP h<sup>2</sup> |
| Matrices | 0.826 | 0.070 | 68.23% | 31.77% | 10.60% | 4.90% | 15.50% |
| Memory | 0.651 | 0.031 | 42.38% | 57.62% | 1.70% | 2.30% | 4.00% |
| RT | 0.308 | 0.026 | 9.49% | 90.51% | 0.70% | 6.70% | 7.40% |
| Symbol Digit | 0.831 | 0.034 | 69.06% | 30.94% | 7.60% | 3.40% | 11.00% |
| Trails-B | 0.976 | 0.035 | 95.26% | 4.74% | 14.20% | 0.70% | 14.90% |
| Tower | 0.853 | 0.080 | 72.76% | 27.24% | 8.30% | 3.10% | 11.40% |
| VNR | 0.717 | 0.024 | 51.41% | 48.59% | 10.90% | 10.30% | 21.20% |
| **Mean %** |  |  | 58.37% | 41.63% | 7.71% | 4.49% | 12.20% |
| Phenotypic g |  |  | Proportion of phenotypic variation explained by phenotypic g and phenotypic s |  |  |  |  |
| Cognitive Trait | Estimate | SE | Common (g) | Specific (s) |  |  |  |
| Matrices | 0.566 | 0.010 | 32.04% | 67.96% |  |  |  |
| Memory | 0.287 | 0.003 | 8.24% | 91.76% |  |  |  |
| RT | 0.361 | 0.003 | 13.03% | 86.97% |  |  |  |
| Symbol Digit | 0.727 | 0.003 | 52.85% | 47.15% |  |  |  |
| Trails-B | 0.819 | 0.003 | 67.08% | 32.92% |  |  |  |
| Tower | 0.545 | 0.010 | 29.70% | 70.30% |  |  |  |
| VNR | 0.519 | 0.004 | 26.94% | 73.06% |  |  |  |
| **Mean %** |  |  | 32.85% | 67.15% |  |  |  |

*Note.* Matrices = Matrix Pattern Completion task; Memory = Memory – Pairs Matching Test; RT = Reaction Time; Symbol Digit = Symbol Digit Substitution Task; Trails-B = Trail Making Test - B; Tower = Tower Rearranging Task; VNR = Verbal Numerical Reasoning Test. All traits are scaled such that higher scores indicate higher cognitive performance. Common (g) R<sup>2</sup> = proportion of genetic variation accounted for genetic g or specific s factors. Specific (s) R<sup>2</sup> = proportion of genetic variance accounted for specific genetic factors associated with each cognitive test. Total SNP heritability = proportion of phenotypic variance accounted for the causal variants identified in each cognitive test univariate GWAS. *Note:* Standardized factor loadings indicate the standardized linear relationship between the factor and each of the cognitive outcomes. Models are fit to LDSC-derived genetic covariance matrices using Genomics SEM. As per best practices for LDSC, genetic covariance matrices were derived using HapMap3 SNPs with minor allele frequencies > 1%, excluding SNPs with INFO < .9 and those from the MHC region.

**Fig. 1.**
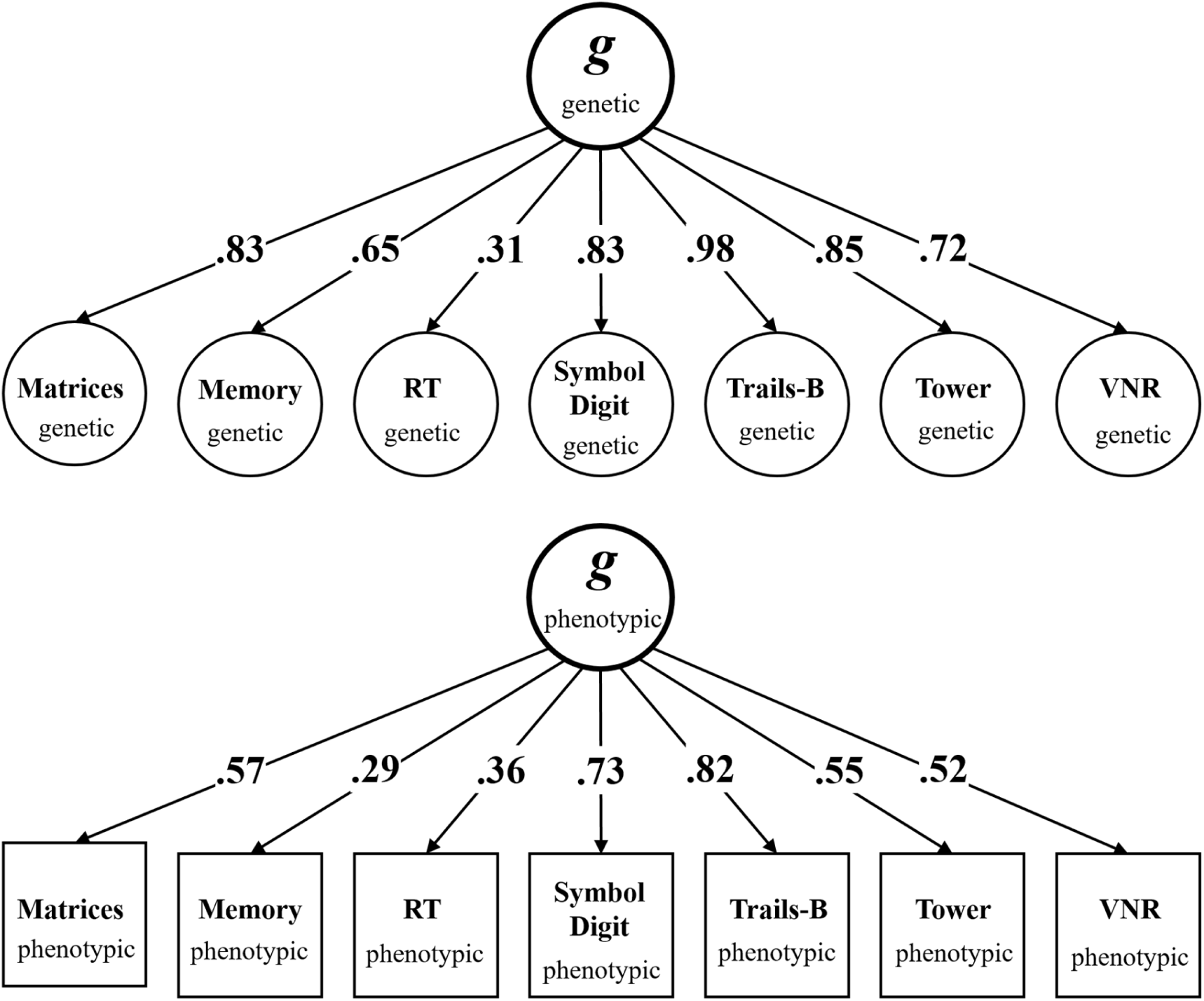
Standardized genetic (top) and phenotypic (bottom) factor solutions for the covariance structure of seven UK Biobank cognitive traits used in the present study. Squares represent observed variables, i.e. the phenotypes that are directly measured. Circles represent latent variables that are statistically inferred from the data, i.e. the genetic and phenotypic *g* factors that are inferred through factor analysis, and the genetic components of the observed phenotypes that are inferred through LD Score Regression. Arrows are standardized factor loadings, which can be interpreted as standardized regression relations with the arrow pointing from the predictor variable to the outcome variable. Genetic factor models were estimated using Genomic SEM (*26*), and phenotypic models were estimated using the lavaan package for R (*41*). Matrix = Matrix Pattern Completion task; Memory = Memory – Pairs Matching Test; RT = Reaction Time; Symbol Digit = Symbol Digit Substitution Task; Trails-B = Trail Making Test – B; Tower = Tower Rearranging Task; VNR = Verbal Numerical Reasoning Test. All variables are scaled such that higher scores indicate better cognitive performance.

We next aimed to determine the contributions of individual genetic loci specifically to genetic *g*, and to distill those from loci associated with other levels of the cognitive hierarchy. We fit a multivariate GWAS of genetic *g* within Genomic SEM (*26*) to distinguish loci relevant to genetic *g* from loci whose patterns of association across the individual traits is inconsistent with their operation on genetic *g*, as indexed by the heterogeneity statistic, Q. We provide detailed explication of the Q statistic and how it can be appropriately interpreted in the *Interpreting the Heterogeneity Statistic* section of the Supplementary Materials. Genome-wide significant loci were defined using FUnctional Mapping and Annotation of genetic associations (FUMA) (*29*); see *Materials and Methods* section of the Supplementary Materials. The GWAS results for genetic *g* and Q are displayed in a Miami plot, Fig. 2. Our method distinguishes four types of significant loci. First, highlighted in red are genome-wide significant loci for genetic *g* that are not genome-wide significant loci for the univariate GWAS analyses of the individual traits. These are discoveries for genetic contributions to differences in general intelligence made by leveraging the joint genetic architecture of the traits. Second, highlighted in blue are genome-wide significant loci for *g* that are also genome-wide significant loci in the univariate GWAS analyses for at least one individual cognitive trait. These hits might otherwise have been confused for loci relevant specifically to the individual trait, when in fact the multivariate results indicate that they are pleiotropic (*23*); i.e. they are relevant to genetic *g*. Third, highlighted in green are genome-wide significant loci for the univariate phenotypes that are not genome-wide significant loci for *g*. These may be loci that are specific to the individual traits, but not genetic *g*. Fourth, highlighted in yellow are loci that evince genome-wide significant heterogeneity (Q), indicating that they show patterns of associations with the cognitive traits that depart from the pattern that would be expected if they were to act on the traits via genetic *g*.

**Fig. 2.**
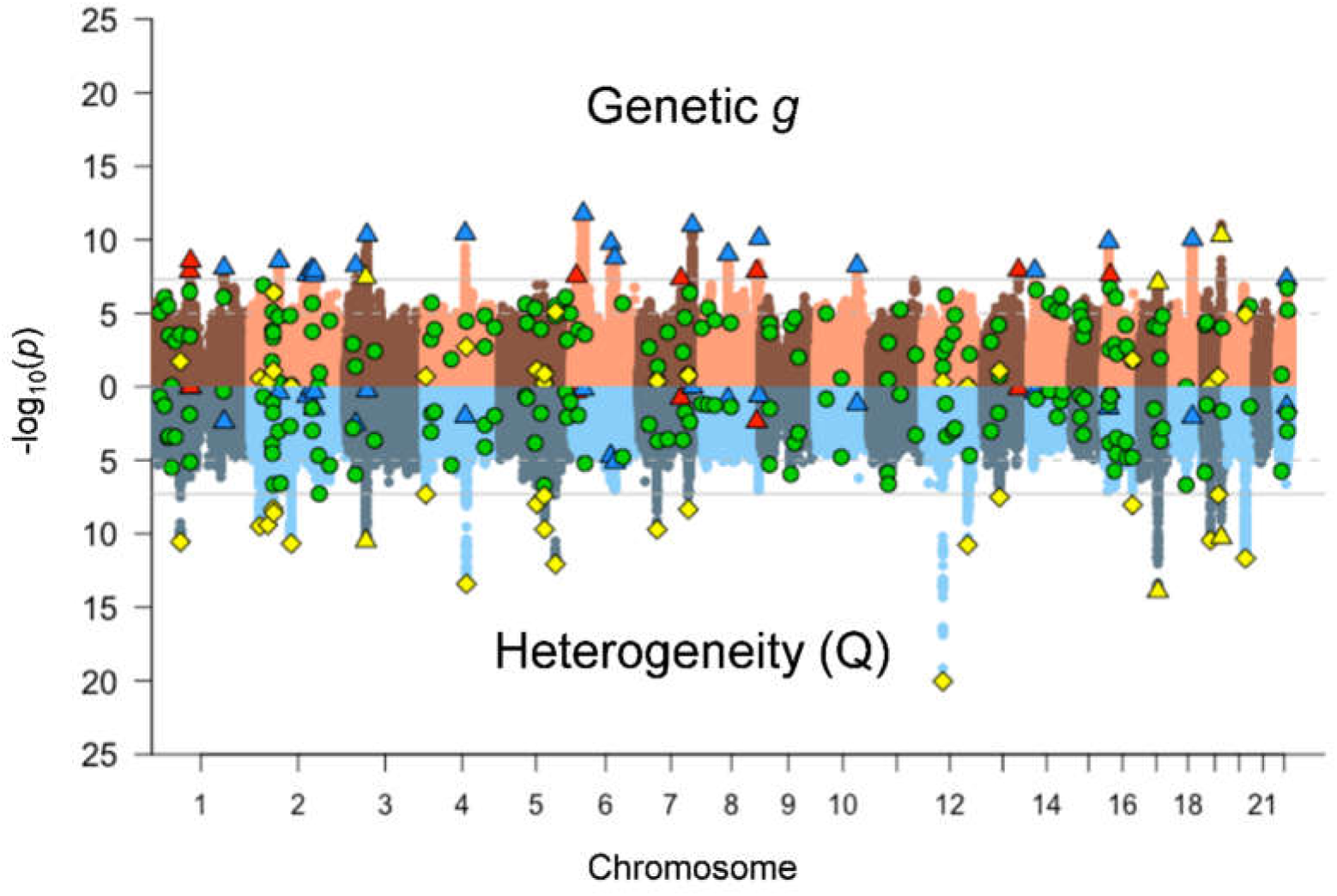
Miami plot of unique, independent hits for genetic *g* (top) and Q (bottom). Q is a heterogeneity statistic that indexes whether a SNP evinces patterns of associations with the cognitive traits that departs from the pattern that would be expected if it were to act on the traits via genetic *g*. The solid grey horizontal lines are the genome-wide significance threshold (p < 5×10^−8^) and the dotted grey horizontal lines are the suggestive threshold (p < 1 ×10^−5^). The following genome wide significant loci are highlighted: *Red triangles* 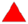: *g* loci unique of univariate loci. *Blue triangles* 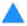: *g* loci in common with univariate loci. *Green circles* 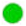: univariate loci not in common with *g* loci. *Yellow triangles* 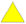: *g* loci in common with Q loci. *Yellow diamonds* 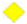: Q loci unique of *g* loci.

Q findings that exceed the genome-wide significance threshold for genetic *g* (yellow triangles) are false discoveries on genetic *g* that are likely driven by a strong signal in a subset of the cognitive traits or in a single cognitive trait. The Q statistic helps to safeguard against these false discoveries. The Q findings that do not surpass the genome-wide significance threshold for genetic *g* (yellow diamonds) are not significantly related to genetic *g* but are significantly heterogeneous in their patterns of associations with the cognitive traits. These loci may be relevant to specific cognitive traits, or to cognitive domains that are intermediate in specificity and generality between *g* and *s*, but not to general intelligence (see Fig. S1). We had coverage, here, of different hierarchically-intermediate traits (e.g. processing speed, memory, reasoning), but they were not measured by multiple tests of each. We were therefore unable formally to model effects on intermediate traits as separate from those on *s* factors specific to the individual cognitive traits.

Inspection of the univariate GWAS results for the individual traits may help to determine the sources of heterogeneity for the Q findings. For instance, a SNP (rs429358) within *APOE*, which is a known risk factor for Alzheimer’s Disease (*30*), was a significant Q finding. With the exception of its association with VNR, this SNP displayed a pattern of associations with the traits that corresponded closely with the degree to which they represented genetic *g*. However, consistent with the inference that *APOE* is specifically relevant for cognitive aging, the SNP displayed a negligible null association with VNR (p=.142), which is a test that shows minimal age-related differences in the UK Biobank data (*31*). Another example of a Q finding is located on Chromosome 17 (chr17: 44021960-44852612), which Davies et al. (*18*) reported to be significantly associated with both general cognitive ability and Reaction Time. From the univariate GWAS results, the largest association for this locus was with Reaction Time, a measure of psychomotor speed with a relatively low loading on genetic *g*. This locus may have a particularly pronounced association with speeded abilities, rather than a general association with genetic *g*. The third Q locus which is also significant for genetic *g* is located on chromosome 3 (chr3:49120040-50234126). This locus has previously-reported associations with general cognitive ability, educational attainment, intelligence, and math ability (*18-20,32*). In the current study, this locus demonstrates significant heterogeneity and displays its largest associations with VNR, Tower, Matrices, and Trails-B, all measures of higher order cognition. It associations with measures of speed and episodic memory (more basic cognitive processes) are negligible.

Overall, we identified 30 genome-wide significant (p < 5×10^−8^) loci for genetic *g*, 23 of which were common with the univariate GWAS of the individual cognitive traits that served as the basis for our multivariate analysis. We identified, in total, 24 genome-wide significant loci for Q, 3 of which were significantly associated with genetic *g* (and therefore likely to be relevant to more specific cognitive traits, and false discoveries on *g*) and 15 of which were significantly associated with at least one individual cognitive trait in the test-specific GWASs.

The importance of these results may be seen by contrasting the results of Trails-B with Reaction Time. For Trails-B, all of the associated loci have already been reported in univariate GWASs of phenotypes that are essentially general cognitive ability (Table 2). On the other hand, most of the Reaction Time loci have not been found in such univariate GWASs of general cognitive ability. Therefore, when identifying loci associated with performance on an individual cognitive test, it is essential to know the extent to which its associations are broadly related to genetic *g* or specifically related to the phenotype under investigation. As we see in the case of Trails-B, none of its related loci associate exclusively to that test (Table 2); they all relate to the superordinate *g*. Failure to take the multivariate structure of the cognitive traits into account may lead to incorrect inferences (*23*)—either that discoveries made in a univariate GWAS of a cognitive trait are generalizable to the broader universe of cognitive traits when they are in fact specific to that trait, or that discoveries made in a univariate GWAS of a cognitive trait are specific to that trait when they are in fact broadly associated with all traits that load on genetic *g.* For instance, although it is not genome-wide significant for any of the univariate GWASs included in the current analysis, our multivariate analysis using Genomic SEM indicates that a locus on chromosome 7 (chr7:104558814-104588161) is associated with genetic *g*. Lee et al. (*32*) have previously reported this locus to be associated with math ability (Table S22), but there are no previously-reported associations with general cognitive function or intelligence. Similarly, we report an association of a locus on chromosome 8 (chr8:64496159-64842662) with Trails-B (an index of executive function, which is itself strongly genetically correlated with *g* (*13*) in the univariate GWAS, and our Genomic SEM analysis indicates that this locus is also related to genetic *g*. Lee et al. (*32*) have previously reported an association between this locus and math ability, but there are no published findings with general cognitive function or intelligence. In both cases, the current results indicate that the loci are broadly relevant to many abilities via genetic *g*, not simply to math ability. Multivariate methods, such as that pioneered here, are necessary in order to distinguish whether a locus is narrowly relevant for an individual cognitive trait or broadly relevant to genetic *g*.

**Table 2.**
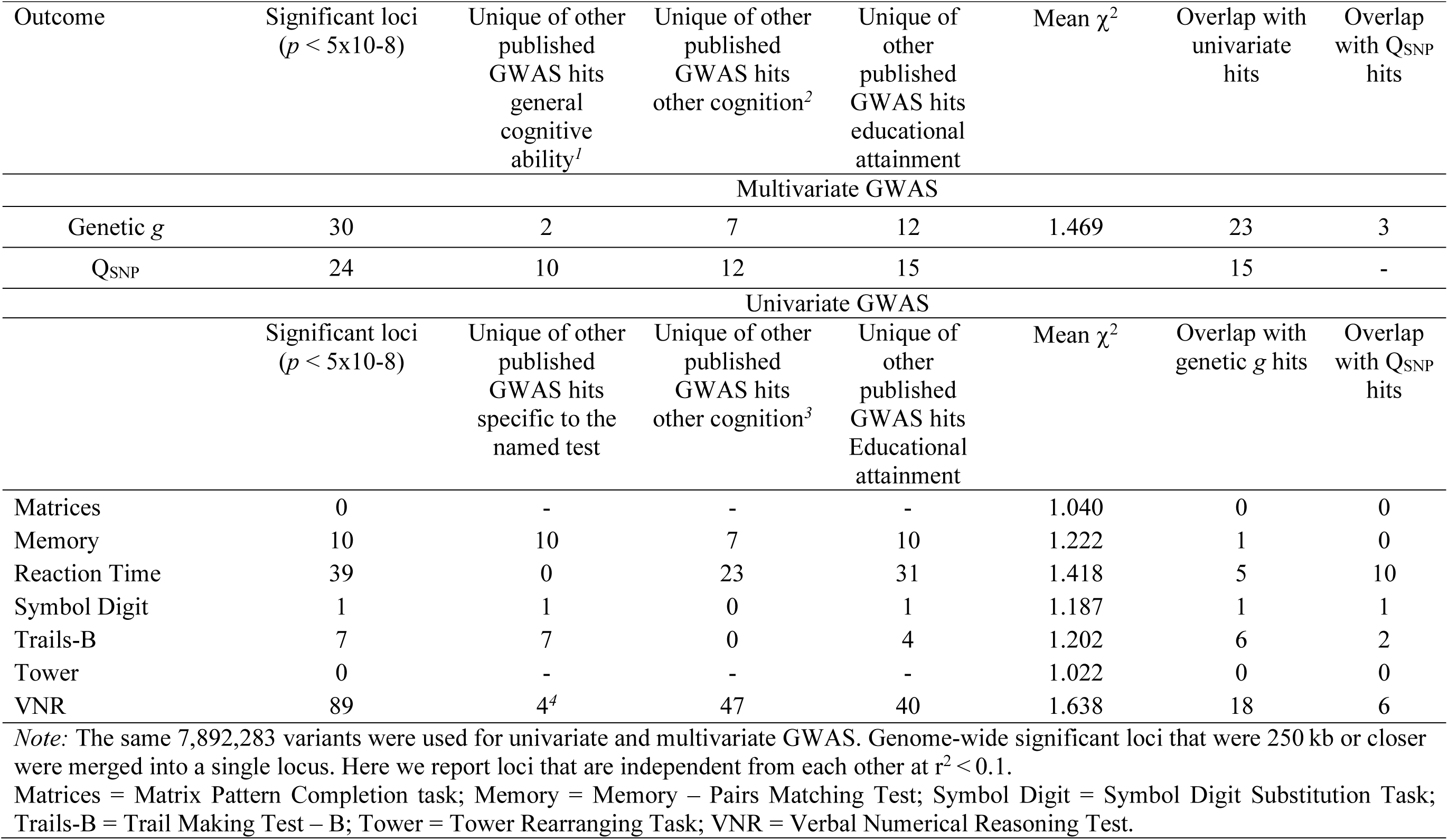
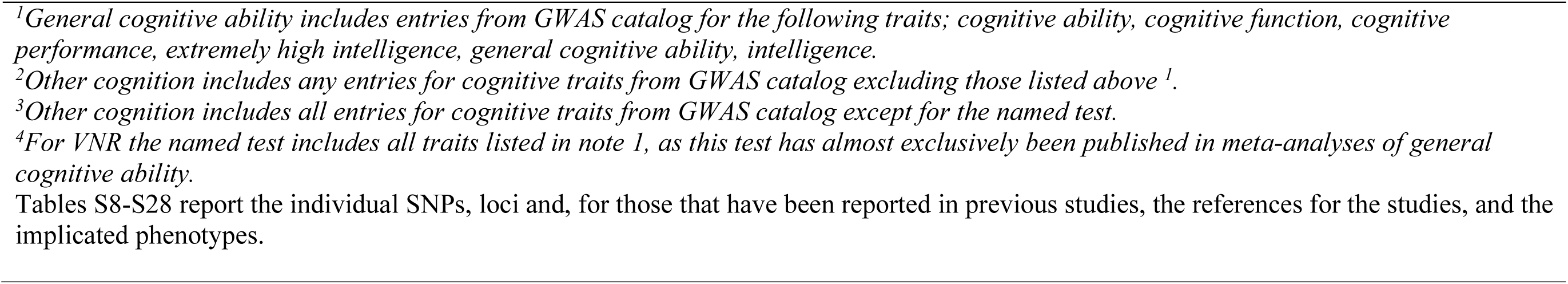
Summary of Multivariate (Genetic *g*) and Univariate GWAS results.

| Outcome | Significant loci<br>($p < 5 \times 10^{-8}$) | Unique of other<br>published<br>GWAS hits<br>general<br>cognitive<br>ability<sup>1</sup> | Unique of other<br>published<br>GWAS hits<br>other cognition<sup>2</sup> | Unique of<br>other<br>published<br>GWAS hits<br>educational<br>attainment | Mean $\chi^2$ | Overlap with<br>univariate<br>hits | Overlap<br>with Q<sub>SNP</sub><br>hits |
| --- | --- | --- | --- | --- | --- | --- | --- |
| Multivariate GWAS |  |  |  |  |  |  |  |
| Genetic *g* | 30 | 2 | 7 | 12 | 1.469 | 23 | 3 |
| Q<sub>SNP</sub> | 24 | 10 | 12 | 15 |  | 15 | - |
| Univariate GWAS |  |  |  |  |  |  |  |
| | Significant loci<br>($p < 5 \times 10^{-8}$) | Unique of other<br>published<br>GWAS hits<br>specific to the<br>named test | Unique of other<br>published<br>GWAS hits<br>other cognition<sup>3</sup> | Unique of<br>other<br>published<br>GWAS hits<br>Educational<br>attainment | Mean $\chi^2$ | Overlap with<br>genetic *g* hits | Overlap<br>with Q<sub>SNP</sub><br>hits |
| Matrices | 0 | - | - | - | 1.040 | 0 | 0 |
| Memory | 10 | 10 | 7 | 10 | 1.222 | 1 | 0 |
| Reaction Time | 39 | 0 | 23 | 31 | 1.418 | 5 | 10 |
| Symbol Digit | 1 | 1 | 0 | 1 | 1.187 | 1 | 1 |
| Trails-B | 7 | 7 | 0 | 4 | 1.202 | 6 | 2 |
| Tower | 0 | - | - | - | 1.022 | 0 | 0 |
| VNR | 89 | 4<sup>4</sup> | 47 | 40 | 1.638 | 18 | 6 |

*Note:* The same 7,892,283 variants were used for univariate and multivariate GWAS. Genome-wide significant loci that were 250 kb or closer were merged into a single locus. Here we report loci that are independent from each other at $r^2 < 0.1$ .
Matrices = Matrix Pattern Completion task; Memory = Memory – Pairs Matching Test; Symbol Digit = Symbol Digit Substitution Task; Trails-B = Trail Making Test – B; Tower = Tower Rearranging Task; VNR = Verbal Numerical Reasoning Test.

Whereas discovery of new genetic associations is not the focus of the current study, we present the largest GWAS to date of a memory phenotype (n = 331,679). From this GWAS, we report seven novel loci, defined as having no previously-reported associations in cognitive GWASs (Table S18). However, some of these loci have been associated with schizophrenia, (*33*) anti-saccade response, (*34*) developmental language disorder (linguistic errors), (*35*) hand grip strength, (*36*) and bone mineral density (*37*) (Table S25).

As expected, the genetic *g* factor identified here displayed strong genetic correlations with general cognitive function from Davies et al., (*18*) (*r*_*g*_ = .90, SE = 0.02), and Savage et al., (*20*) (2018) (*r*_*g*_ = .87, SE = 0.05), which were univariate GWASs of broad cognitive phenotypes, and that of Hill et al., (*19*) (*r*_*g*_ = .80, SE = 0.02), which was a GWAS of intelligence that incorporated educational attainment GWAS summary statistics to boost power via multi-trait analysis of GWAS (MTAG) (*38*). We emphasize, however, that it is not relevant to compare the numbers of genetic loci found to be related to cognitive traits in the present study with those previous studies. Rather, the focus here has been the novel contribution of distilling *g*-specific from non-*g*-related genetic loci. However, we have described particularly illustrative examples of loci found to be related to *g* and to specific traits, and all associations are comprehensively reported in the Supplementary Materials.

Negative genetic correlations were found between our genetic *g* and Alzheimer’s disease (*r*_*g*_ = -.34, SE = 0.05), Schizophrenia (*r*_*g*_ = -.37, SE = 0.03), and ADHD (*r*_*g*_ = -.23, SE = 0.04). Additionally, genetic *g* had significant positive genetic associations with total Brain Volume (*r*_*g*_ = .20, SE = 0.04), and Longevity (*r*_*g*_ = .26, SE = 0.03). Genetic *g* had a positive genetic correlation with Educational Attainment (*32*) (*r*_*g*_ =.48, SE = 0.02) that is lower than those found between Education and previous GWASs of cognitive ability (all estimated at r > .69) (*18–20*). To determine whether this lower association was driven by the inclusion of speeded measures as indicators of genetic *g*, we re-estimated the genetic correlations using a genetic *g* factored formed from the cognitive traits that excluded speeded measures (RT, Trails-B, and Symbol Digit). This version of the genetic *g* factor accounted for 69.7% (SE = 11%) of the genetic variance in the four remaining cognitive traits. This version of genetic *g* produced a somewhat higher genetic correlation between genetic *g* and Educational Attainment (*r*_*g*_ =.60) that continued to be lower than those found between Educational Attainment and previous GWASs of general cognitive ability. We suspect that previous GWASs of general cognitive ability might have tapped more academic forms of cognitive function (i.e. crystallized abilities, such as verbal knowledge) than those tapped by the constellation to cognitive tests (even when excluding the speeded tests) used to form genetic *g* in the current report. Table S6 reports genetic correlations of genetic *g* and other major intelligence GWASs with educational attainment, neural phenotypes, and longevity.

We next created polygenic scores (PGSs) for genetic *g* and the individual UK Biobank cognitive traits and used them to individually and simultaneously predict variance in cognitive performance and educational attainment in an independent study, Generation Scotland; N=6,950 unrelated individuals (Table S7). The cognitive measures were Wechsler Logical Memory (episodic memory), Mill Hill Vocabulary (crystallized knowledge), Wechsler Digit Symbol Substitution (processing speed), and Verbal Fluency (semantic fluency), as described previously (*39,40*). We also calculated the first unrotated principal component of the cognitive measures to index phenotypic *g*, and the first unrotated principal component of the cognitive measures excluding Mill Hill Vocabulary to index a more fluid *g.* Consistent with the Genomic SEM findings that individual cognitive outcomes are associated with a combination of genetic *g* and specific genetic factors, we observed a pattern in which many of the regression models that included both the polygenic score (PGS) from genetic *g* and test-specific PGSs were considerably more predictive of the cognitive phenotypes in Generation Scotland than regression models that included only either a genetic *g* PGS or a PGS for a single test. A particularly relevant exception involved the Digit Symbol Substitution test in Generation Scotland, which is a similar test to the Symbol Digit Substitution test in UK Biobank, for which we derived a PGS. We found that the proportional increase in R^2^ in Digit Symbol by the Symbol Digit PGS beyond the genetic *g* PGS was <1%, whereas the genetic *g* PGS improved polygenic prediction beyond the Symbol Digit PGS by over 100%, reflecting the power advantage obtained from integrating GWAS data from multiple genetically correlated cognitive traits using a genetic *g* model. An interesting counterpoint is the PGS for the VNR test, which is unique in the UK Biobank cognitive test battery in indexing verbal knowledge (*24,31*). Highlighting the role of domain-specific factors, a regression model that included this PGS and the genetic *g* PGS provided substantial incremental prediction relative to the genetic *g* PGS alone for those Generation Scotland phenotypes most directly related to verbal knowledge: Mill Hill Vocabulary (62.45% increase) and Educational Attainment (72.59%).

## Discussion

Until now, research on the positive manifold of correlations among cognitive traits has been phenotypic in nature, or has made inferences regarding the roles of genes only indirectly, using twin approaches. Therefore, we have previously not known to what extent genetic factors underlie the hierarchy of cognitive variation. Here, we have identified a genetic *g* factor using molecular genetic data, and we have discerned loci that operate on genetic *g* from those that operate on more specific cognitive traits. Harking back to Spearman’s 1904 paper’s title, the present study has contributed to understanding how measured “general intelligence” is “determined”.

## Acknowledgements

This article benefited from valuable discussions with Michel Nivard.

## Author Contributions

I.J.D. and E.M.T-D. jointly conceived of the idea, designed the study, and formulated the analytic plan. J. F. and G. D. performed the analyses, with contributions from A.D.G. I.J.D. and E.M.T-D. wrote the paper, with contributions from J. F. and G. D. All authors contributed to editing the paper.

## Data Availability

Complete summary GWAS results from this paper will be made available at the time of publication at https://www.ccace.ed.ac.uk/node/335. Raw data for UK Biobank can be requested at https://www.ukbiobank.ac.uk/register-apply/. Raw data for Generation Scotland can be requested at https://www.ed.ac.uk/generation-scotland/using-resources/access-to-resources/access-process. Code to perform common factor modeling and multivariate GWAS within Genomic SEM can be found at https://github.com/MichelNivard/GenomicSEM/wiki.

## Supplementary Material

### Method

#### Sample

Data from the UK Biobank study was used for the present study (https://www.ukbiobank.ac.uk/). The UK Biobank is a biomedical prospective cohort study, which collected a wide range of genetic and health related measures from a national sample of community dwelling participants from the UK. Ethical approval for the UK Biobank was granted from the Research Ethics Committee (11/NW/0382). This study uses European ancestry genome-wide genotyped data from seven cognitive tests with varying sample sizes across phenotypes.

#### Cognitive Tests

##### Reaction Time (*n* = 330,024)

This test was self-administered by participants at the baseline UK Biobank assessment. In this task, pairs of either identical or different cards were presented on a computer screen. If the two cards were identical, participants had to push a button as quickly as possible. Reaction time (RT) score corresponded with the time, in milliseconds, to identify the matching cards in four trials. Participants were presented with 12 trials in total. The first five trails were used as a practice. Of the remaining seven trials, four presented identical cards. The score is the mean time, in milliseconds, for these four trials. Whereas there are only a few trials, internal consistency is good (Cronbach α = 0.85).

##### Matrix Pattern Recognition (*n* = 11,356)

The non-verbal fluid reasoning Matrix Pattern Recognition test is an adaptation of the Matrices test included in the COGNITO battery (*42*), which is similar to the well-known raven’s Progressive Matrices test. This test was self-administered during the assessment centre imaging visit. This test involves inspection of an abstract grid pattern with a piece missing in the lower right-hand corner. The pattern has a logical order. The participant is asked to select the correct multiple-choice option at the bottom of the screen to complete the logical pattern both horizontally and vertically. This 15-item test aims at assessing the ability to solve non-verbal, non-numerical problems using novel and abstract materials. The score is the total number of correctly solved items in three minutes.

##### Verbal Numerical Reasoning (*n* = 171,304)

At the baseline assessment center visit, a sub-sample of UK Biobank participants self-administered the verbal-numerical reasoning test. Participants were asked 13 multiple-choice questions that assessed verbal and numerical problem solving. The score was the number of questions answered correctly in two minutes. This test has been shown to have adequate test-retest reliability (r = 0.65) (*43*) and internal consistency (Cronbach α = 0.62) (*31*). The verbal-numerical reasoning test was also administered to three sub-samples of participants at the first repeat assessment visit, the assessment center imaging visit, and during the web-based cognitive assessment. In the web-based version of this test there was an additional question, thus the maximum score was 14. In the current analysis the verbal numerical reasoning score used is from the first testing occasion for each participant.

##### Symbol Digit Substitution (*n* = 87,741)

The symbol digit substitution test was self-administered during both the assessment center imaging visit and the web-based cognitive assessment. Participants were shown a key, pairing shapes with numbers. Participants were asked to use the key to fill the maximum number of empty boxes with the corresponding number paired with shapes in a series of rows. The score is the number of correct symbol-digit matches made in 60 seconds. Those with a score coded as 0 and those with a score greater than 70 had their score set to missing. In this analysis, the scores used were from the first testing occasion for each participant.

##### Memory – Pairs Matching Test (*n* = 331,679)

At the baseline UK Biobank assessment, memory was measured using a ‘pairs matching’ task. In this self-administered task, participants are shown a randomly arranged, four by three grid of 12 ‘cards’, with six pairs of matching symbols, for five seconds. The symbols were then hidden, and the participant was instructed to select, from memory, the locations of the pairs that matched, in the fewest possible number of attempts. There was no time limit for this task. The memory score was the total number of errors made during this task before all pairs were identified.

##### Tower rearranging (*n* = 11,263)

This test was self-administered during the imaging assessment center visit. It is similar to the well-known ‘Tower of Hanoi’ task. Participants were presented with a display (display A) containing three different colored hoops arranged on three pegs (towers). Another display (display B) was shown underneath display A, with the three hoops arranged differently. The task involves deciding how many moves it would take to change display A into display B. The score was the number of correctly-completed trials achieved in three minutes.

##### Trail Making Test – B (*n* = 78,547)

This test is a computerized version of the Halstead-Reitan Trail Making Test (*44*). The trail making test was self-administered during both the assessment center imaging visit and the web-based cognitive assessment. In part B of the test, participants were presented with the numbers 1-13, and the letters A-L arranged quasi-randomly on a computer screen. The participants were instructed to switch between touching the numbers in ascending order, and the letters in alphabetical order (e.g., 1-A-2-B-3-C) as quickly as possible. The score was the time (in seconds) taken to successfully complete the test. Those with a score coded as 0 (denoting “Trail not completed”) had their score set to missing. In this analysis, the scores used were from the first testing occasion for each participant.

#### Genotyping

Prior to release of the UK Biobank genetic dataset, QC measures were applied; these are described in Bycroft et al (*45*). We applied additional quality control filters prior to analyses; individuals were removed sequentially based on non-British ancestry, high missingness, high relatedness (samples which have more than 10 putative third-degree relatives), and gender mismatch. Our analysis sample included 332,050 unrelated participants of European descent with high-quality genotyping. 80,639,280 autosomal single nucleotide polymorphism (SNP) variants were analyzed (imputation reference panels included UK10K haplotype, 1000 Genomes Phase 3, and Haplotype Reference Consortium (HRC) panels); all variants had a minor allele frequency ≥ 0.000009 and an imputation quality score of ≥ 0.1.

#### Genome-wide association analyses

Univariate genome–wide association analyses were performed for each UK Biobank cognitive phenotype using a linear association test in BGENIE (*45*). All cognitive phenotype scores were residualized for age, assessment center (where required), genotype batch, array, and 40 genetic principal components prior to analysis. For phenotypes which were collected across multiple testing occasions, a separate GWAS was performed for each occasion and these were meta-analyzed using METAL (*46*). As the size of the subsets of individuals for each phenotype vary greatly, an additional QC filter to remove SNPs with a minor allele count < 25 was applied to all GWAS summary results prior to further analyses.

#### Factor Models

We performed genetic factor analysis on the UK Biobank cognitive phenotypes with Genomic SEM (*26*) and phenotypic factor analysis with the lavaan package for R (*41*). In both genetic and phenotypic factor analysis, a common factor model specifies that *k* phenotypes are described as linear functions of a smaller set of *m* (continuous) latent variables: **y** = **Λη+ε**. In this equation, **y**is a *k* × 1 vector of indicators, ε is a *k* ×; 1 vector of residuals, **η** is an *m*×;1 vector of common factors, and Λ is a *k* ×; *m* matrix of factor loadings, i.e. regressions relating the common factors to the set of indicators. In the genetic factor model, y represents the genetic components of the GWAS phenotypes, whereas, in the phenotypic factor model, y represents the phenotypes themselves. The model-implied covariance matrix of a CFA is **Σ**(*θ*) = = **ΛΨΛ′**+**Θ**, where **Ψ** is an *m* × *m* latent variable covariance matrix (in the case of a single common factor, **Ψ** is simply equal to the variance of the factor, which we fix to 1 for scaling identification purposes), and **Θ** is a *k* ×; *k* matrix of covariances among the residuals, ε (typically a diagonal matrix, to indicate that all indicator residuals are assumed to be independent of one another). A set of parameters (*θ*) is estimated such that the fit function indexing the discrepancy between the model-implied covariance matrix, ∑(θ), and the empirical covariance matrix, *S*, is minimized.

For genetic factor modelling in Genomic SEM (*26*), S is a genetic covariance matrix that is empirically estimated in stage prior to model-fitting using a multivariable extension of Linkage Disequilibrium Score Regression (LDSC) (*25*). For phenotypic factor modelling, S is a phenotypic covariance matrix that is empirically estimated from the raw phenotypic data. The fit function used to estimate the model parameters takes into account the precision of the elements of the S matrix, along with their sampling dependencies (which are needed to appropriately account for sample overlap across the GWAS phenotypes) in the form of a sampling covariance matrix, V, that is estimated using a jackknife resampling procedure in the multivariable extension of LDSC available in Genomic SEM. Model fit is considered good when ∑(θ) closely approximates S. For Genomic SEM, the fit function used was Diagonally Weighted Least Squares, with a sandwich estimator. For Phenotypic modeling, the fit function used was maximum likelihood.

In Genomic SEM, goodness-of-fit of the model is assessed by means of the standardized root mean square residual (SRMR), model χ^2^, Akaike Information Criterion (AIC), and Comparative Fit Index (CFI). For phenotypic factor modelling, we additionally consider Root Mean Square Error of Approximation (RMSEA). Hu and Bentler (*47*) have proposed the following criteria for a good fit: Comparative Fit Index (CFI) > 0.95; Tucker-Lewis Index (TLI) > 0.95; Root Mean Squared Error of Approximation (RMSEA) < 0.08.

#### Estimation of SNP-based heritability and genetic correlations using GCTA-GREML

Our primary means of estimating the UK Biobank’s cognitive phenotypes’ SNP-based heritability and genetic correlations was with the multivariable version of LDSC available in Genomic SEM, as described above. However, in order to verify that the estimated genetic correlation matrix was consistent across estimation methods relying on different assumptions, we additionally implemented GCTA-GREML to estimate SNP-based heritability (*28*) and genetic correlations (*27*). Due to computational requirements for the bivariate GCTA-GREML analyses a subset of individuals was created and used for all of the GCTA-GREML analyses. This subset was created by performing listwise deletions for reaction time, memory, VNR, symbol digit substitution, and TMT-B; n = 72,583. The same covariates were included in all GCTA-GREML analyses as for the SNP-based association analyses. One individual was excluded from any pair of individuals who had an estimated coefficient of relatedness of >0.05 to ensure that effects due to shared environment were not included.

#### Genetic correlations with neural phenotypes and longevity

We extended the factor models in Genomic SEM to estimate the genetic correlations between the genetic *g* factor from the UK Biobank cognitive phenotypes and each of nine collateral phenotypes in turn: Educational Attainment (*32*); general cognitive function from Davies et al. (*18*), Savage et al. (*20*)^15^,, and Hill et al. (*19*)^16^; total brain volume from UKB (*48*)^17^; Alzheimer’s disease (*49*)^18^, Schizophrenia (*50*)^19^, Attention Deficit Hyperactive Disorder (ADHD) (*50*)^19^, Autism Spectrum Disorder (ASD) (*50*)^19^, and longevity (*51*)^20^.

#### Multivariate GWAS in Genomic SEM

Genomic SEM (*26*) was used to conduct a multivariate GWAS of the seven UK Biobank cognitive phenotypes, with the genetic *g* factor as the GWAS target. As the typical unit variance scaling cannot be directly specified in a model in which the latent factor is a dependent variable, we specified unit loading scaling (with Matrices as the reference indicator; Table S32). Summary statistics for the individual tests were restricted to SNPs with an MAF > 1%, an INFO score > 0.6, and to SNPs that were present for all seven cognitive tests. The summary statistics were also filtered to SNPs present in the European only 1000 Genomes Phase 3 reference panel, as the SNP minor allele frequencies from the reference panel are necessary to obtain their variances for inclusion in the genetic covariance (*S*) matrix. Using these QC steps, 7,892,283 SNPs were present across all seven cognitive tests.

#### Genome-wide significant loci characterization using FUMA

Genome-wide significant loci were defined from the SNP-based association results, using FUnctional Mapping and Annotation of genetic associations (FUMA) (*29*). The SNP2GENE function was used to identify independent significant SNPs defined as SNPs with a *P*-value of ≤5 × 10^−8^ and independent of other genome wide significant SNPs at r^2^ < 0.6. Tagged SNPs, for use in subsequent annotations, were then identified as all SNPs that had a MAF ≥ 0.0005 and were in LD of r^2^ ≥ 0.6 with at least one of the independent significant SNPs. These tagged SNPs included those from the 1000 genomes reference panel and need not have been included in the GWAS performed in the current study. Genome-wide significant loci that were 250 kb or closer were merged into a single locus. Lead SNPs were defined as independent significant SNPs that were independent from each other at r^2^ < 0.1. We performed look-ups on all tagged SNPs (r^2^ > 0.6) within each locus, including all 1000 genomes SNPs; previously reported genome-wide significant findings are detailed in Tables S22-28.

#### Functional annotation implemented in FUMA

Only SNPs reported in novel genomic loci, defined as those loci which do not have any previously reported cognitive or educational attainment associations, identified for Q, VNR, and memory were annotated for functional consequences on gene functions using ANNOVAR (*52*) and the Ensembl genes build 85. A CADD score (*53*), RegulomeDB score (*54*), and 15-core chromatin states (*55–57*) were obtained for each SNP. Functionally-annotated SNPs were then mapped to genes based on physical position on the genome. Intergenic SNPs were mapped to the two closest up- and down-stream genes which can result in their being assigned to multiple genes. (Tables S29-S31)

#### Polygenic prediction

Generation Scotland: the Scottish Family Health Study (GS) is a family-structured, population-based cohort study recruited between 2006 and 2011. Participant recruitment occurred in Glasgow, Tayside, Ayrshire, Arran, and North-East Scotland, yielding a total sample size of 24,084 with an age range between 18 and 100 years, and up to four generations per family, of which we selected one participant per family. A full cohort description is provided elsewhere (*39,40*) and online at http://www.generationscotland.org/. Ethical approval for GS was obtained from the Tayside Committee on Medical Research Ethics (on behalf of the National Health Service). Genotyping, using the Illumina HumanOmniExpressExome-8 v1.0 chip, was performed at the Edinburgh Clinical Research Facility, University of Edinburgh (*58*). Participants were removed from GS if they had contributed to both GS and UK Biobank (n = 622). For the PGS analyses 6,950 unrelated GS participants were retained.

The cognitive measures available in GS were Wechsler Logical Memory (episodic memory), Mill Hill Vocabulary (crystallized knowledge), Wechsler Digit Symbol Substitution (processing speed), and Verbal Fluency (semantic fluency), as described previously (*39,40*). We created a phenotypic g using the first unrotated principal component of the cognitive measures, and also a fluid g using the first unrotated principal component of the cognitive measures excluding the Mill Hill Vocabulary scores.

Polygenic profile scores were created using PRSice version 2 (https://github.com/choishingwan/PRSice) (*59*). Summary results from the genetic g and univariate cognitive test GWAS were used to create polygenic profile scores for the GS individuals. Prior to creating these scores SNPs with a MAF < 0.01 were removed and clumping was used to obtain SNPs in linkage disequilibrium with an r2 < 0.25 within a 250 kb window. Polygenic profile scores for the individual cognitive tests were created using all available SNPs from the GWAS summary results. For genetic g, all SNPs located within significant Q loci were removed from the GWA summary results prior to the profile scores being created.

Linear regression models were used to examine the associations between the polygenic profile scores and cognitive performance and educational attainment in GS. All models included age at measurement, sex, and 10 genetic principal components to adjust for population stratification. We created regression models fitting each polygenic score individually, a multivariate model including all eight polygenic scores (genetic g, reaction time, memory, matrix, symbol digit substitution, trail making test B, and tower rearranging) and, a series of models which fitted the genetic g polygenic score plus one individual cognitive test score. From these models we were able to determine contributions of genetic g and each individual UK Biobank cognitive test to prediction of variance in cognitive performance and educational attainment in an independent sample, GS.

### Interpreting the Heterogeneity Statistic

Here we provide a description of what the Genomic SEM heterogeneity statistic (Q) indexes, and how we can appropriately interpret the detected heterogeneity by investigating the univariate GWAS results for the individual phenotypes.

Q indexes the extent to which model misfit occurs for a *common pathway* model in which the effects of a given SNP on the individual phenotypes are specified to occur exclusively via a single effect of the SNP on the latent factor (Fig. S9, left panel) compared to a less restrictive *independent pathways model* in which the effects of a given SNP on the individual phenotypes are specified to occur directly on those phenotypes (Fig. S9, right panel). In other words, low Q indicates that the SNP plausibly acts on the latent factor, whereas high Q indicates that the SNP does not plausibly act on the latent factor.

Under the *common pathway* model, the expected SNP effects 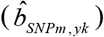 on phenotype *k*, is *b*_*SNPm,F*_ × *λ*_*k*_, i.e.:

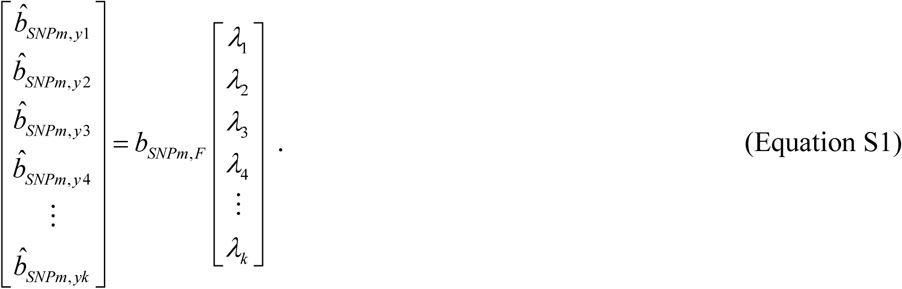

Misfit occurs when the vector of expected SNP effects on phenotypes 1 through *k* deviates from the vector of observed SNP effects on phenotypes 1 though *k* (*b*_*SNPm*, *yk*_), as estimated from the univariate GWASs. If the effects of the SNP on the individual phenotypes occur exclusively by way of the effect of the SNP on the common factor, the vector of observed SNP effects should be proportional to the vector of unstandardized loadings of those phenotypes on the common factor, and Q will be low. If, however, the vector of observed SNP effects <u>is not</u> proportional to the vector of unstandardized loadings of those phenotypes on the common factor, Q will be high, and a model in which the SNP effects on the phenotypes occur exclusively via the common factor will be rejected.

When Q is high for a given SNP, the linear association between the vector of univariate regression coefficients, *b*_*SNPm*, *yk*_, and the vector of unstandardized factor loadings, λ_*k*_, will be weaker, and there may be one or more outliers. Note that we <u>do not</u> necessarily expect that a SNP that acts exclusively on the common factor to have relatively equal univariate associations with each phenotype (this will only occur if the factor loadings are all relatively similar). Rather, if the SNP acts exclusively on the common factor, we expect the univariate associations to scale with the unstandardized factor loadings for the corresponding phenotypes. For instance consider a phenotype with a relatively low unstandardized factor loading. We expect that a SNP that acts directly and exclusively on the common factor to have a relatively lower association with that phenotype compared to its associations with the other phenotypes. In fact, if the association with that phenotype is comparable to the association of that SNP with other phenotypes, Q will be high.

We next explain why it is the SNP’s beta coefficients, and not its Z statistics or p values, that must be explored to investigate heterogeneity. Importantly, when the different phenotypes differ dramatically in their sample sizes, SNP heritabilities, or polygenicity, the Z statistics (or p value, which is derived from the Z statistic) for a given SNP and each phenotype are *not* expected to be proportional to the magnitude of the unstandardized factor loadings for those phenotypes. For instance, imagine a scenario in which Q is 0 (no heterogeneity) for a particularly SNP, such that the correlation between the vectors of betas and factor loading is 1.0. We would still likely see differences in Z statistics (and p values) across phenotypes that do not correspond with their unstandardized factor loadings. All else being equal, the phenotypes with the largest Ns will have very high Z statistics and those with the smallest Ns will have very low Z statistics (and may not be significant). If we investigate the Z statistics or p values, we may incorrectly infer that the SNP is relevant to the high N phenotypes but not the low N phenotypes. However, if we investigate the betas and rely on the Q statistic, we will come to a very different (and more correct) conclusion. If Q is 0 (such that the method of correlated vectors produces r=1.0), and the SNP effect on the common is genome-wide significant, we will correctly conclude that we have identified a SNP that plausibly acts on the phenotypes via the factor.

Now consider what happens when Q is high for a particular SNP. We may be interested in identifying the source(s) of the heterogeneity across phenotypes. The same principles as above hold; we must investigate the betas. If we investigate the Z statistics or p values, we will simply conclude that the SNP is specific to the phenotypes for which the univariate GWASs are more highly powered, whether this is true or not.

Next, we consider how these principles apply to the results with respect to genetic *g*. Table S33 provides the univariate GWAS summary statistics for the lead SNPs from the 3 loci that are genome-wide significant for both Q and genetic *g*.

The first hit considered is for a lead SNP within the APOE gene, which is a well-known risk factor for Alzheimer’s Disease. Fig. S10 is the scatterplot of the betas against the unstandardized factor loadings. It can be seen that the betas correspond very closely to the factor loadings for all traits except VNR, which is a test that does not decline with age. The betas for memory and RT are also low, but these are traits with relatively lower unstandardized loadings, so these observations are unlikely to contribute directly to high Q. VNR is an outlier because its beta is low relative to its factor loading. Note that only Symbol Digit and Trails-B pass the genome-wide significance threshold, so one (likely incorrect) interpretation would be that this is a SNP that is only relevant to those two traits. However, remember that Q is examining heterogeneity in betas across all traits (not just the significant ones), and whether they scale with factor loadings. If our goal was to tally the intersection of univariate hits for the same SNPs across traits, we would not need multivariate methods. However, our goal is to evaluate how these SNPs operate within a formal multivariate model. Importantly, Q is genome-wide significant for this SNP, which is why we investigate it further. However, differences in the significance of the SNP associations for the individual traits are not directly relevant for interpreting the genome-wide significant Q statistic.

The second hit considered is for a lead SNP within a locus on Chromosome 17. Fig. S11 is the scatterplot of the betas against the factor loadings. It can be seen that there is not much correspondence between the factor loadings and the SNP effects, even with outliers removed. The SNP has its strongest association with RT (a measure of psychomotor speed), but it also has a sizable association with Symbol Digit. The RT association is the only genome-wide significant univariate association for this SNP, but one hesitates to conclude that this is a SNP that is specific to RT, given the magnitude of the Digit Symbol beta. A more conservative conclusion would be that this is a SNP that is more broadly related to speeded abilities.

The last hit considered is for a lead SNP within a locus on Chromosome 3. Fig. S12 is the scatterplot of the betas against the factor loadings. First, it is important to observe that although there appears to be good correspondence between the factor loadings and the betas, this isn’t exactly the case, as two of the betas are slightly negative. Because the factor loadings are all positive, it is not possible for a vector that contains both positive and negative SNP effects to be proportional to the factor loadings. Rather, there appears to be two clusters of SNP effects. One cluster (Memory, RT, and Digit Symbol; all tests of basic mental processes) is characterized by associations that are very close to 0. A second cluster (Trails-B, Tower, Matrices, VNR; all tests of higher order cognition) is characterized by similarly sized positive associations. This SNP only exhibits Z statistics surpassing the suggestive threshold (*p*<1×10^−5^) for Trails-B and VNR, but in fact the SNP’s beta coefficient for Tower is larger (.029) than its coefficient for Trails-B (.023). Tower (*n* = 11,263) is simply less well powered than is Trails-B (*n* = 78,547). One would be very hesitant to say that this SNP is only relevant for Trails–B and VNR as its regression relations with Tower and Matrices are very similar in magnitude as those for Trails–B and VNR. One sensible interpretation of this pattern is that that this SNP is relevant for higher-order cognition, but not basic cognitive processes.

Some points of caution are important to keep in mind. First, the formal hypothesis being tested, for which a genome-wide multiple testing correction is made, is the omnibus test of heterogeneity (Q). Interpretation of the specific pattern of SNP-phenotype associations following identification of genome-wide significant Q loci is post-hoc, and should therefore be considered tentative. Nevertheless, for the reasons described above, basing such investigation of the individual phenotype associations on regression coefficients is more appropriate than basing such investigations on Z-statistics or p-values. This is because Genomic SEM is a formal framework for modeling effect sizes across traits, and is not simply a method of pooling p values. Importantly, interpreting genome-wide significant Q loci in terms of on p values or Z statistics from disproportionally powered univariate GWASs can lead to interpretations that fail to account for the fact that “the difference between significant and not significant is [not necessarily] itself significant”. (*60*)

Second, overfitting is always bound to be a problem when SNPs are identified on the basis of surpassing a stringent significance threshold for their associations. In conventional univariate GWAS it is well-known that the effect sizes for genome-wide significant SNPs are likely to be overestimated in the discovery sample. In multivariate GWAS, such as the analyses conducted on genetic *g*, when SNPs are identified on the basis of surpassing a stringent threshold for heterogeneity, there is likely to be collider bias that builds in artifactual dependencies between individual SNP effect sizes. Short of having well-powered independent validation data to re-estimate individual SNP effects for Q hits, this collider bias will be difficult to fully resolve, and must be considered when making interpretations.

---

**Fig. S1.**
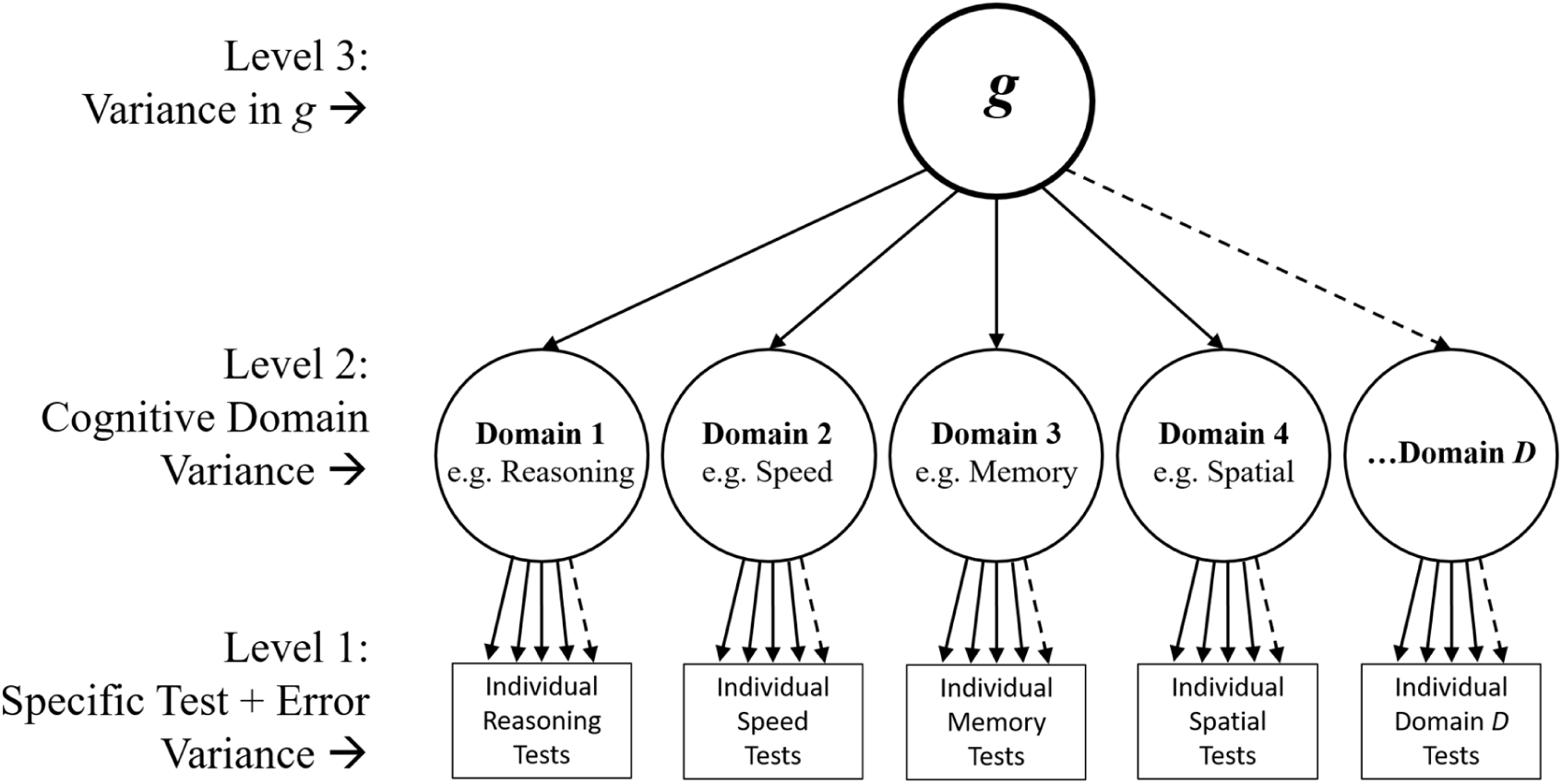
Hierarchical structure of intelligence differences (after Deary (*61*) & Carroll (*5*). Cognitive abilities composing intelligence are measured via a variety of diverse cognitive tests (Level 1). Spearman (*4,62*) discovered that person-to-person differences in performance on many different cognitive tests are moderately positively correlated. Later work refined this discovery (*5,63–66*), and articulated the hierarchical model, after observing particularly strong positive correlations among tests within cognitive domains (Level 2), such that latent traits representing the domains of performance can be extracted to represent their common variance. People who have strengths in one domain also tend to have strengths in other domains, such that a general intelligence factor, *g*, can be extracted (Level 3). This hierarchical structure of intelligence differences is well-established (*5*). Approximately 40% of the variation in performance on the individual tests of a diverse cognitive battery is accounted for by *g*, approximately 25% of additional variation is accounted for by the cognitive domains after taking *g* into account, and then approximately 35% is explained by factors that are specific to the individual tests and by measurement error. When only one test per domain is available, the sources of variation stemming from levels 1 and 2 to cannot be separated, but *g* can still be extracted. It has been found that, so long as the tests used to measure cognitive abilities are sufficiently diverse, almost the same *g* factor is always extracted (6,*62*).

**Fig. S2.**
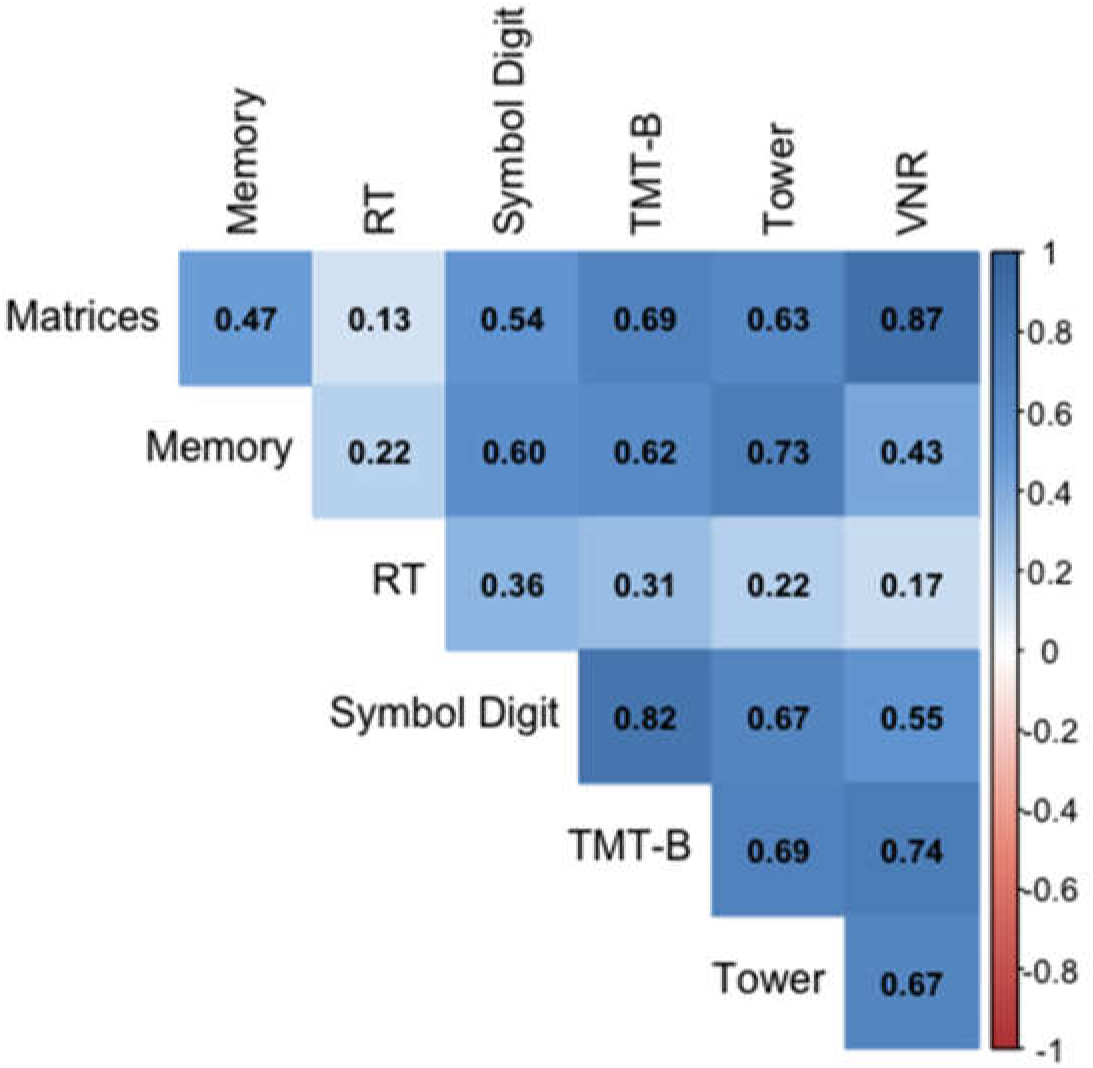
Heat-map of LDSC-estimated genetic correlations among the seven UK Biobank cognitive phenotypes. Matrix = Matrix Pattern Completion task; Memory = Memory – Pairs Matching Test; RT = Reaction Time; Symbol Digit = Symbol Digit Substitution Task; Trails-B = Trail Making Test − B; Tower = Tower Rearranging Task; VNR = Verbal Numerical Reasoning Test.

**Fig. S3.**
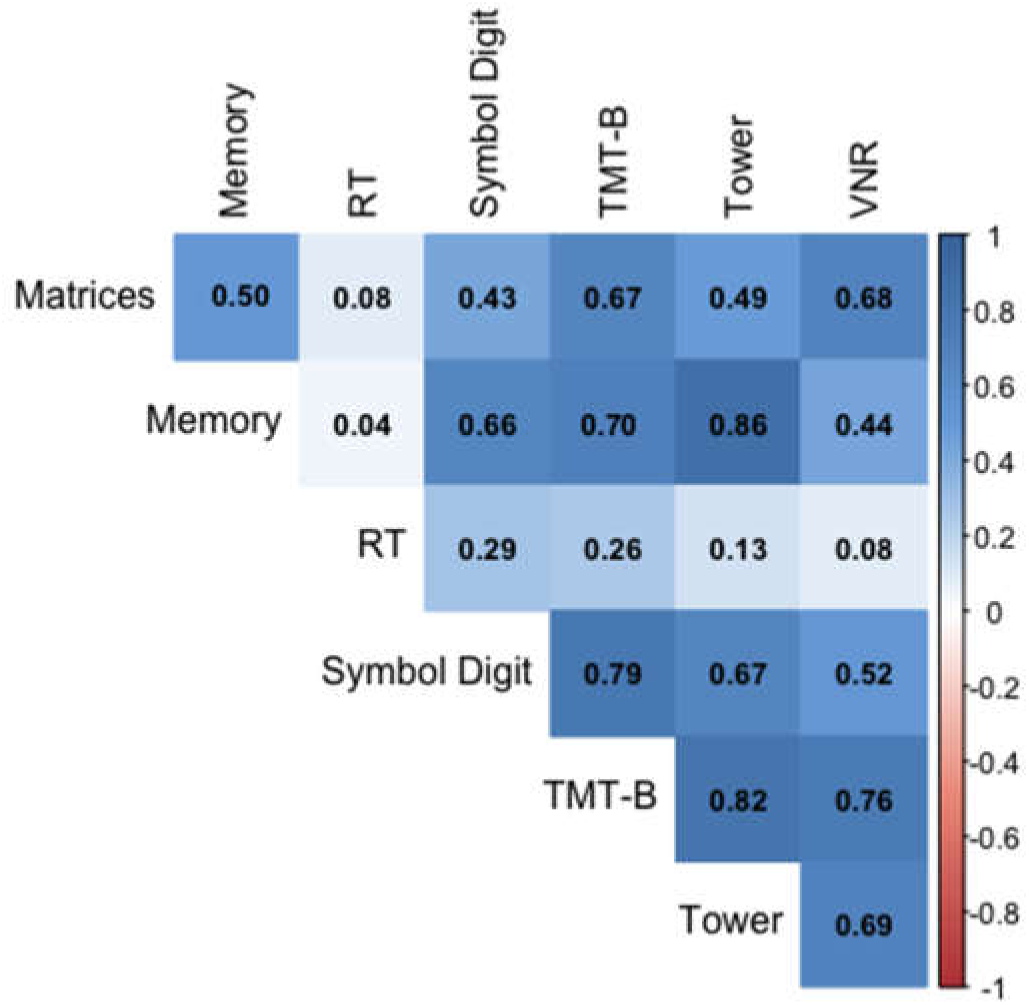
Heat-map of GREML-estimated genetic correlations among the UK Biobank cognitive phenotypes. Matrix = Matrix Pattern Completion task; Memory = Memory – Pairs Matching Test; RT = Reaction Time; Symbol Digit = Symbol Digit Substitution Task; Trails-B = Trail Making Test − B; Tower = Tower Rearranging Task; VNR = Verbal Numerical Reasoning Test.

**Fig. S4.**
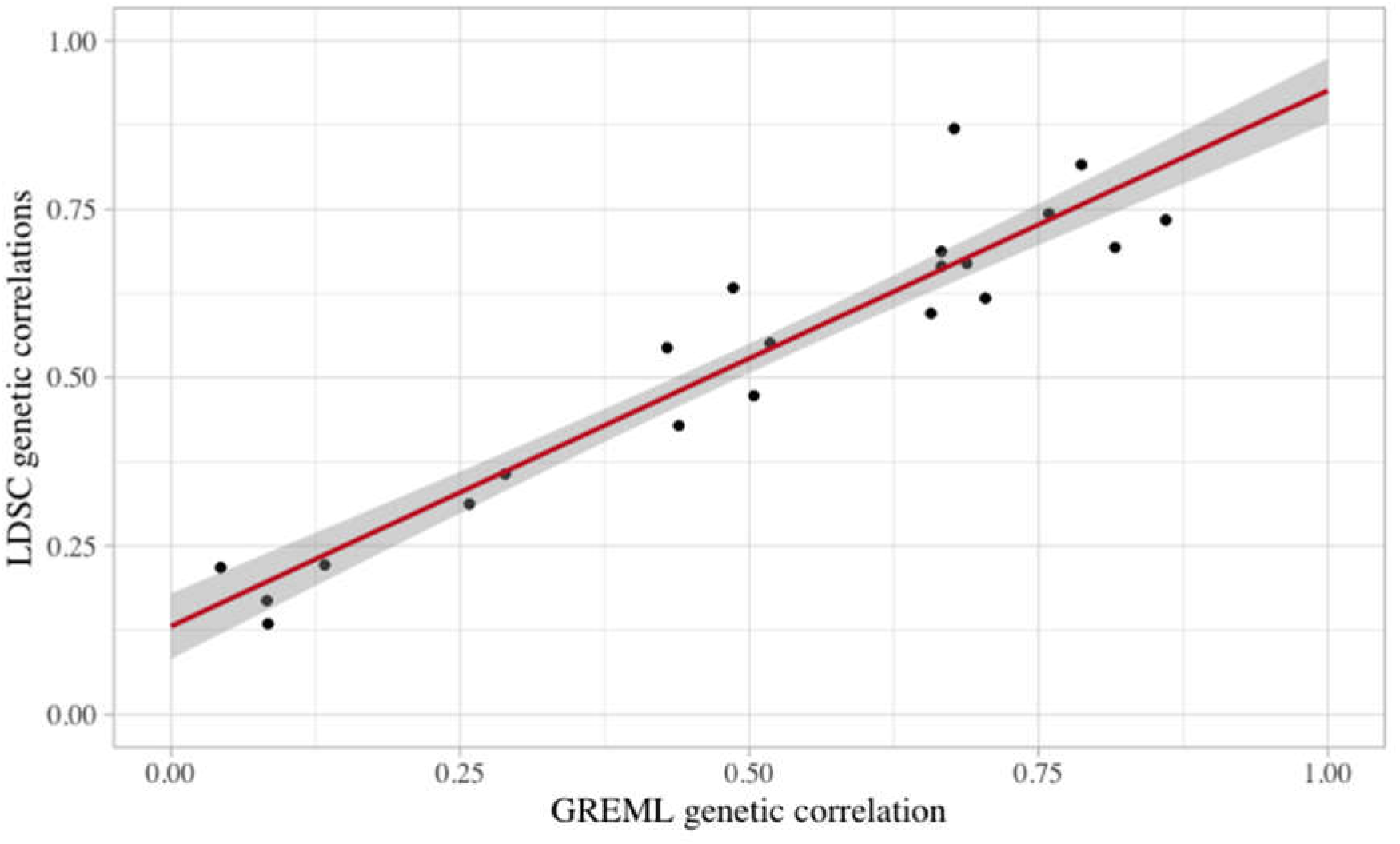
Scatterplot of LDSC and GREML genetic correlations (from Figs S2 and S3) among UK Biobank cognitive phenotypes. Note: Shaded area represents 95% confidence interval.

**Fig. S5.**
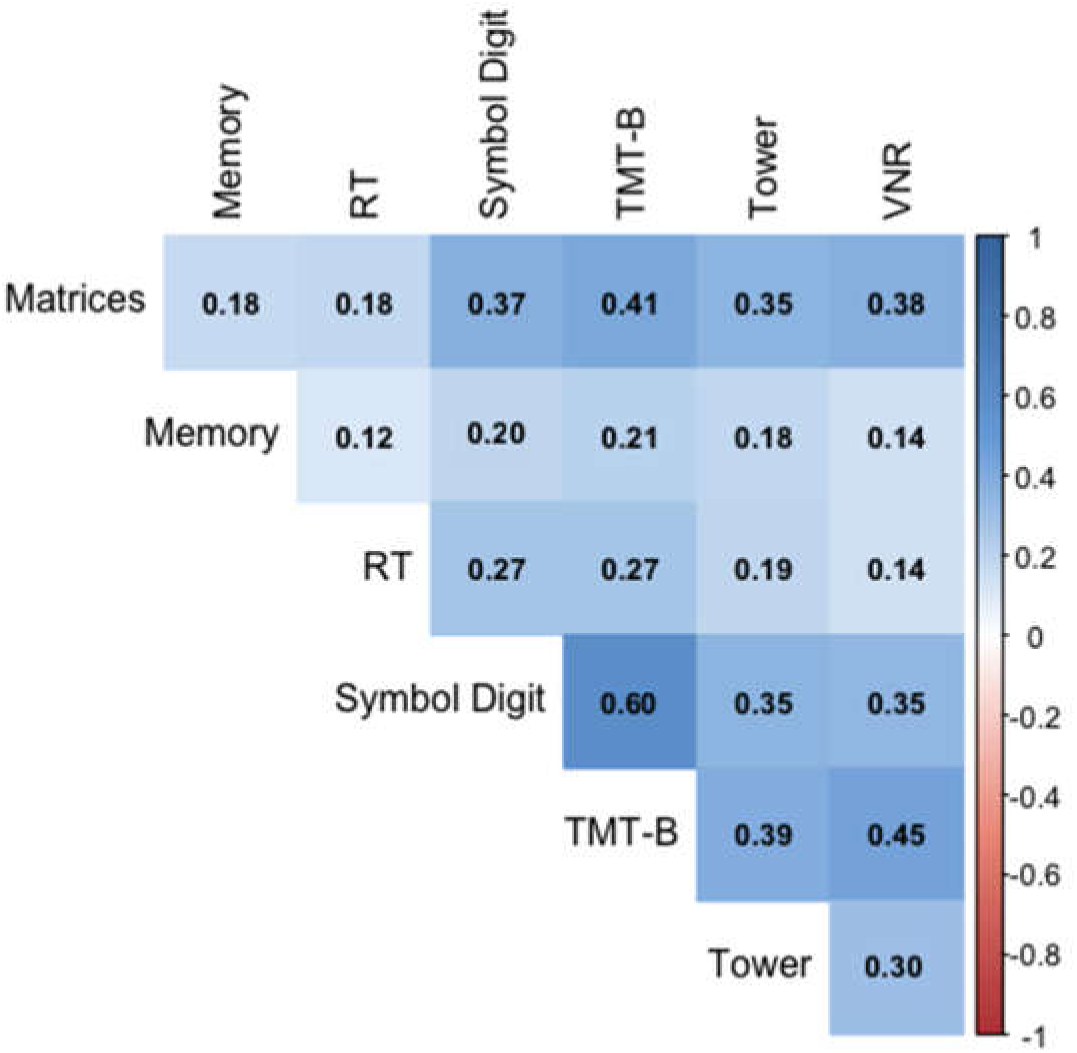
Heat-map of phenotypic correlations among the seven UK Biobank cognitive tests. Matrix = Matrix Pattern Completion task; Memory = Memory – Pairs Matching Test; RT = Reaction Time; Symbol Digit = Symbol Digit Substitution Task; Trails-B = Trail Making Test – B; Tower = Tower Rearranging Task; VNR = Verbal Numerical Reasoning Test.

**Fig. S6.**
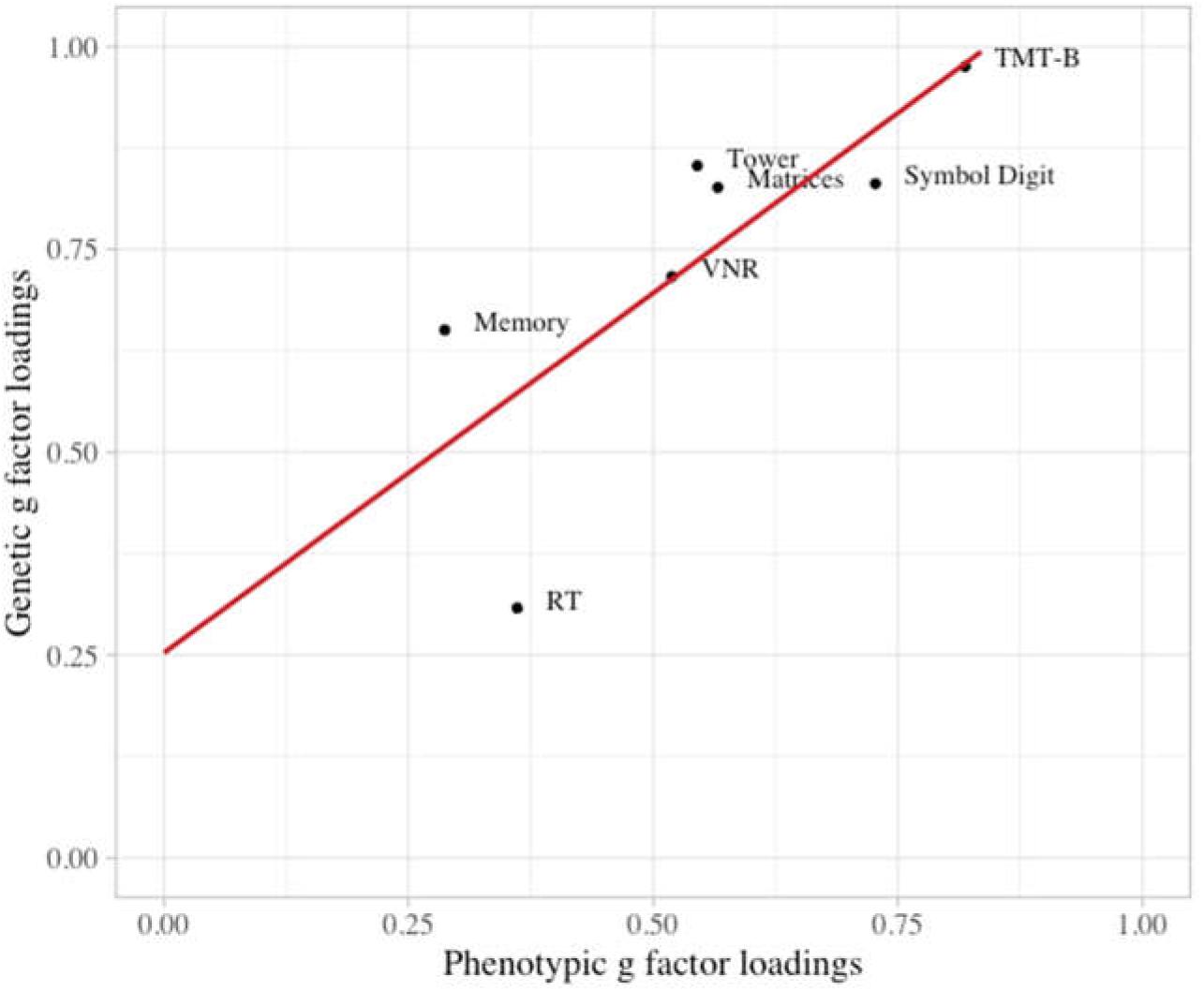
Scatterplot of phenotypic and genetic *g* factor loadings. Matrix = Matrix Pattern Completion task; Memory = Memory – Pairs Matching Test; RT = Reaction Time; Symbol Digit = Symbol Digit Substitution Task; Trails-B = Trail Making Test – B; Tower = Tower Rearranging Task; VNR = Verbal Numerical Reasoning Test. Note: the regression intercept of the regression line displayed in this figure is .253, and the unstandardized slope is. 887. The intercept of .253 indicates somewhat higher genetic than phenotypic factor loadings, and the slope of. 887 indicates close correspondence between their orderings.

**Fig. S7.**
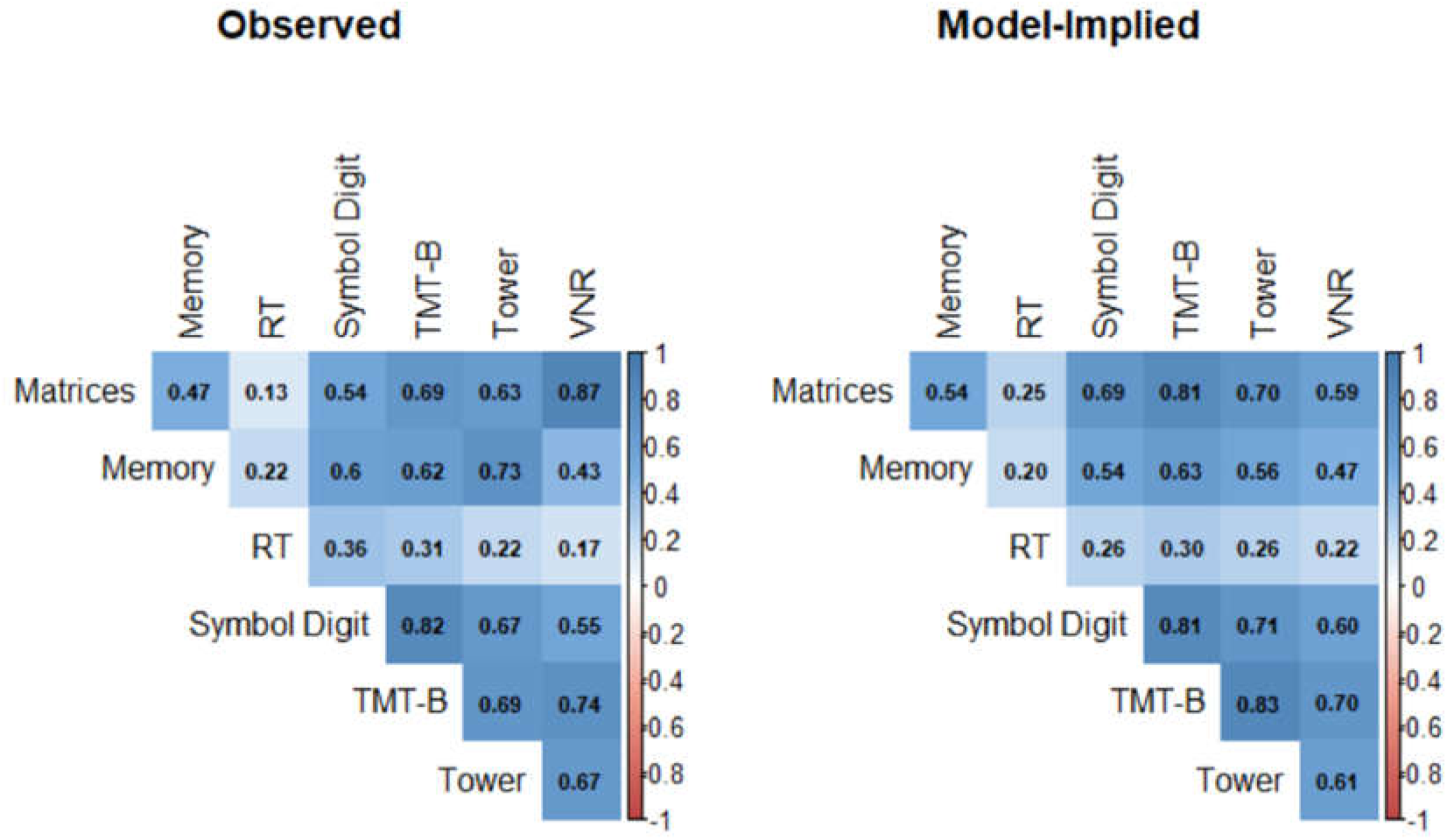
Heat-map of LDSC-estimated (observed) and genetic *g* model-implied genetic correlations among UK Biobank cognitive phenotypes. Matrix = Matrix Pattern Completion task; Memory = Memory – Pairs Matching Test; RT = Reaction Time; Symbol Digit = Symbol Digit Substitution Task; Trails-B = Trail Making Test – B; Tower = Tower Rearranging Task; VNR = Verbal Numerical Reasoning Test.

**Fig. S8.**
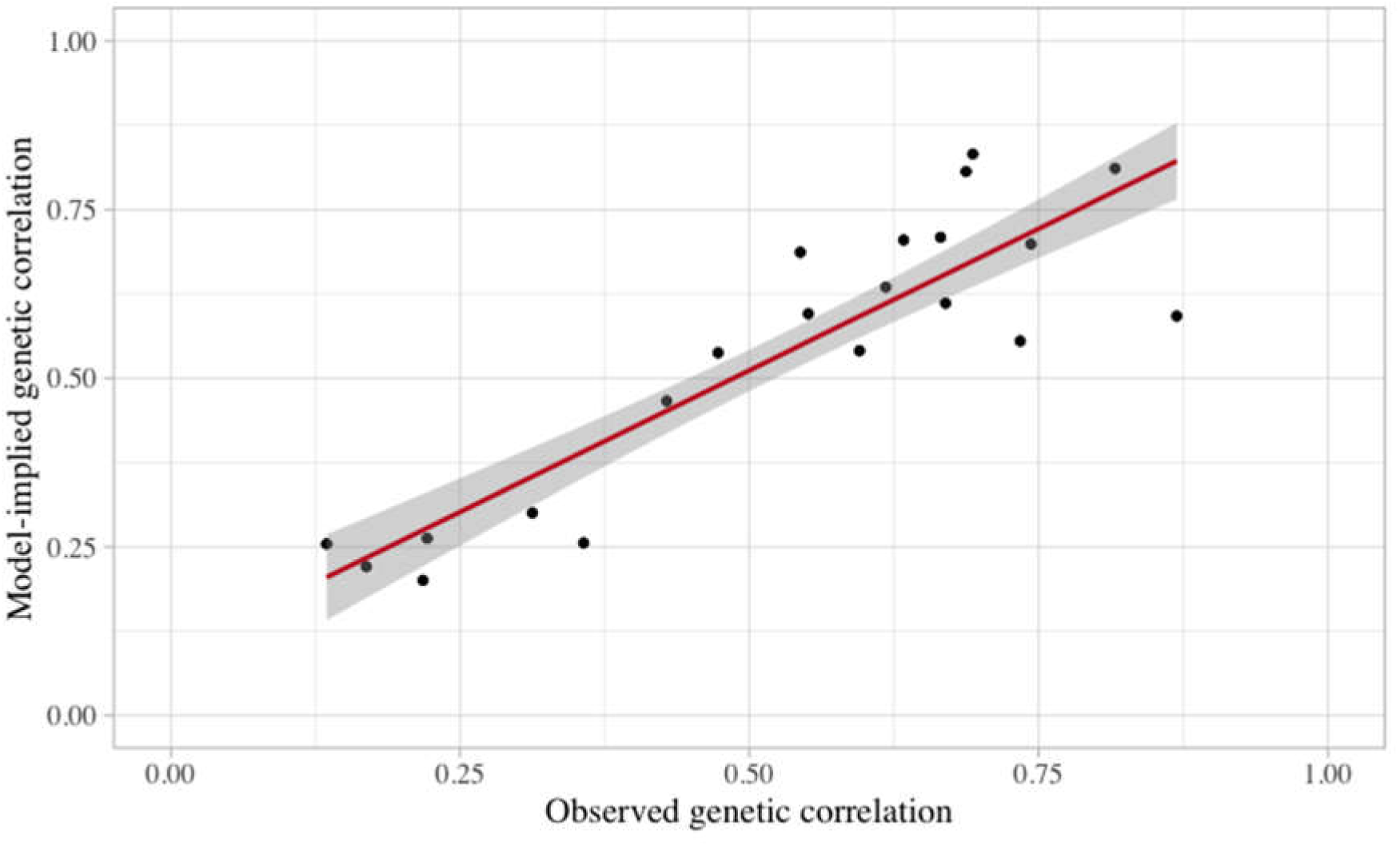
Scatterplot of observed and model-implied genetic correlations (from Fig. S5) among UK Biobank cognitive phenotypes. Note: Shaded area represents 95% confidence interval.

**Fig. S9.**
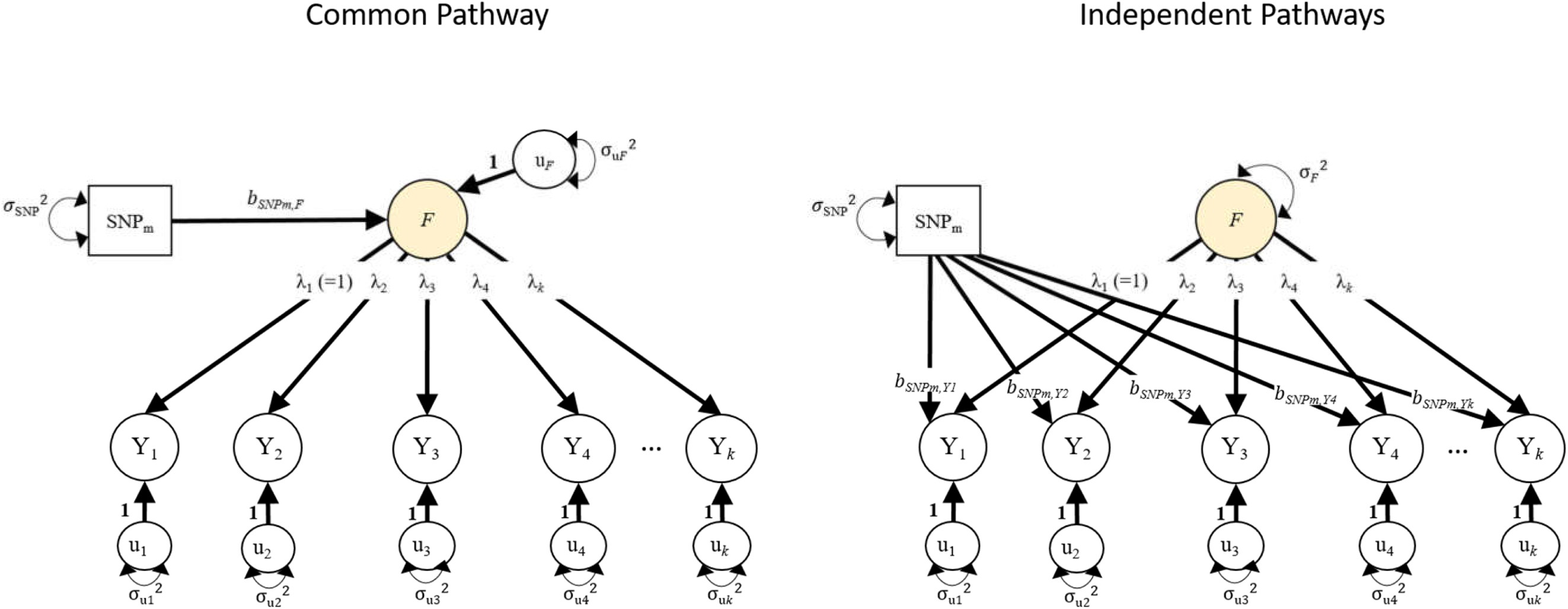
Unstandardized path diagrams for *common pathway* (left) and *independent pathways* (right) models used to compute the Genomic SEM heterogeneity statistic (Q) for a multivariate GWAS of a single common factor. In this example, F is a common genetic factor of the genetic components of *k* GWAS phenotypes (Y_1_-Y_*k*_). Each model is run once for each SNP, *m*. Single-headed arrows are regression relations, and double-headed arrows are variances. Paths labeled 1 are fixed to 1 for model identification purposes. All other paths represent freely estimated model parameters. Q represents the decrement in model fit of the *common pathway* model relative to the more restrictive *independent pathways* model. Q is a χ^2^ distributed test statistic with k-1 degrees of freedom, representing the difference between the *k* SNP-phenotype *b* coefficients in the independent pathways model and the 1 SNP-factor *b* coefficient in the *common pathway* model.

**Fig. S10.**
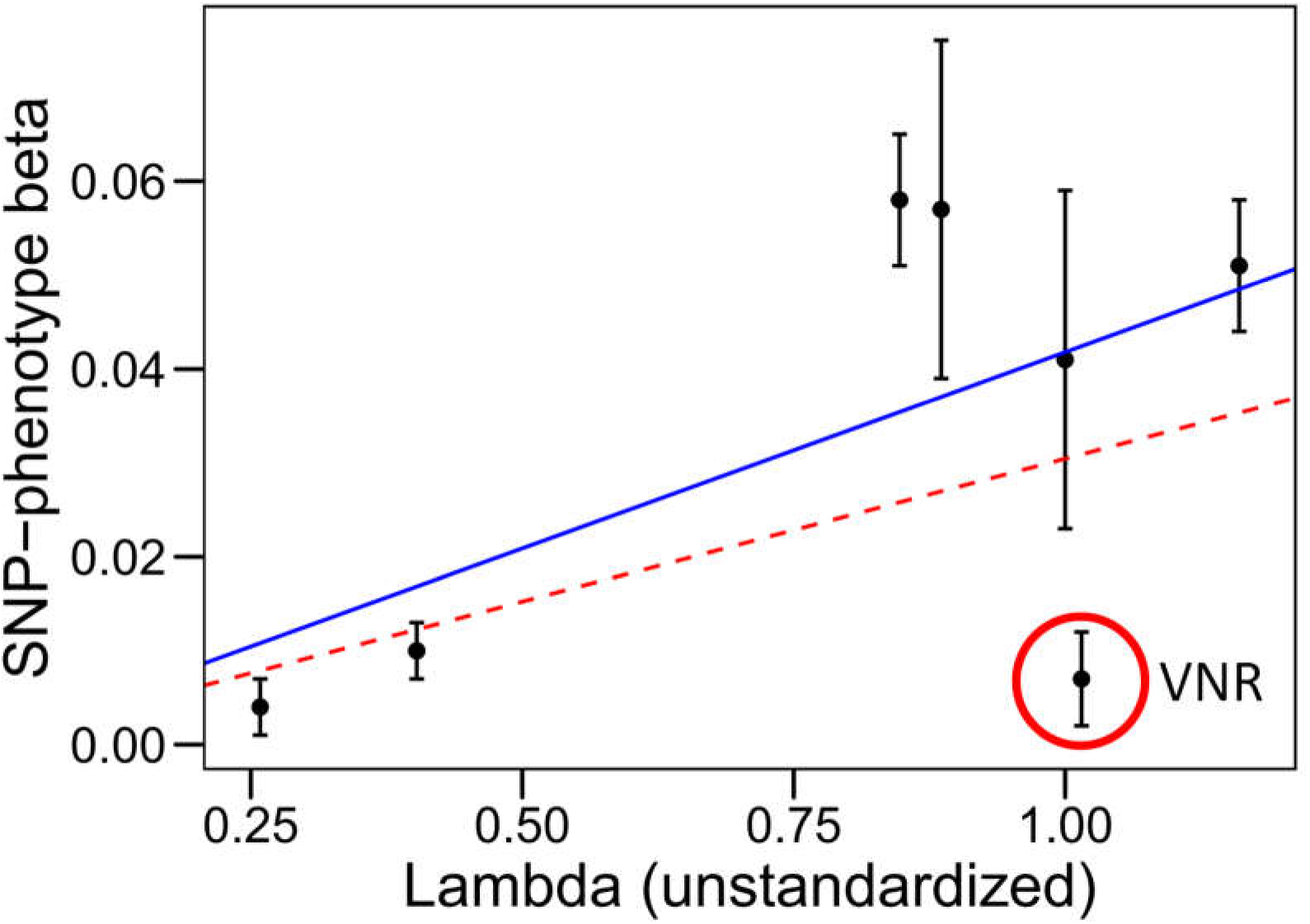
Scatter plot of SNP-phenotype regression coefficients (betas) against unstandardized genetic factor loadings for the associated phenotypes, for lead SNP **rs429358** within the APOE gene. Error bars represent standard errors of the SNP-phenotype betas. The dashed red line represents the regression line based on all seven data points. The solid blue line represents the regression line after excluding VNR. In order to correspond directly with Equation S1, both regression lines were estimated with their intercepts fixed to zero, using weights equal to the inverse of the squared standard errors of the betas.

**Fig. S11.**
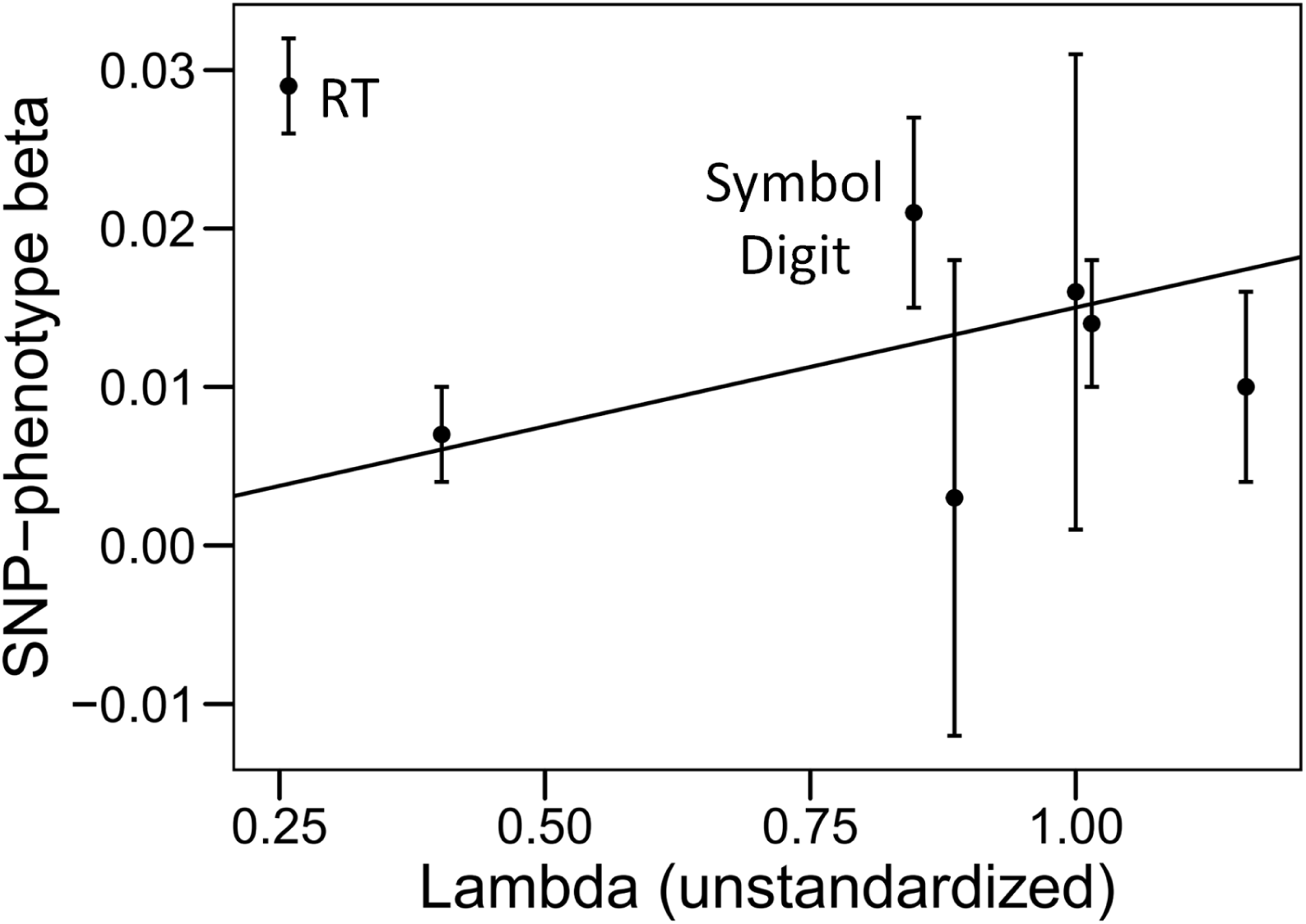
Scatter plot of SNP-phenotype regression coefficients against unstandardized genetic factor loadings for the associated phenotypes, for **rs273534** within a locus on Chromosome 17. Error bars represent standard errors of the SNP-phenotype betas. The solid black line represents the regression line based on all seven data points. In order to correspond directly with Equation S1, the regression line was estimated with its intercept fixed to zero, using weights equal to the inverse of the squared standard errors of the betas.

**Fig. S12.**
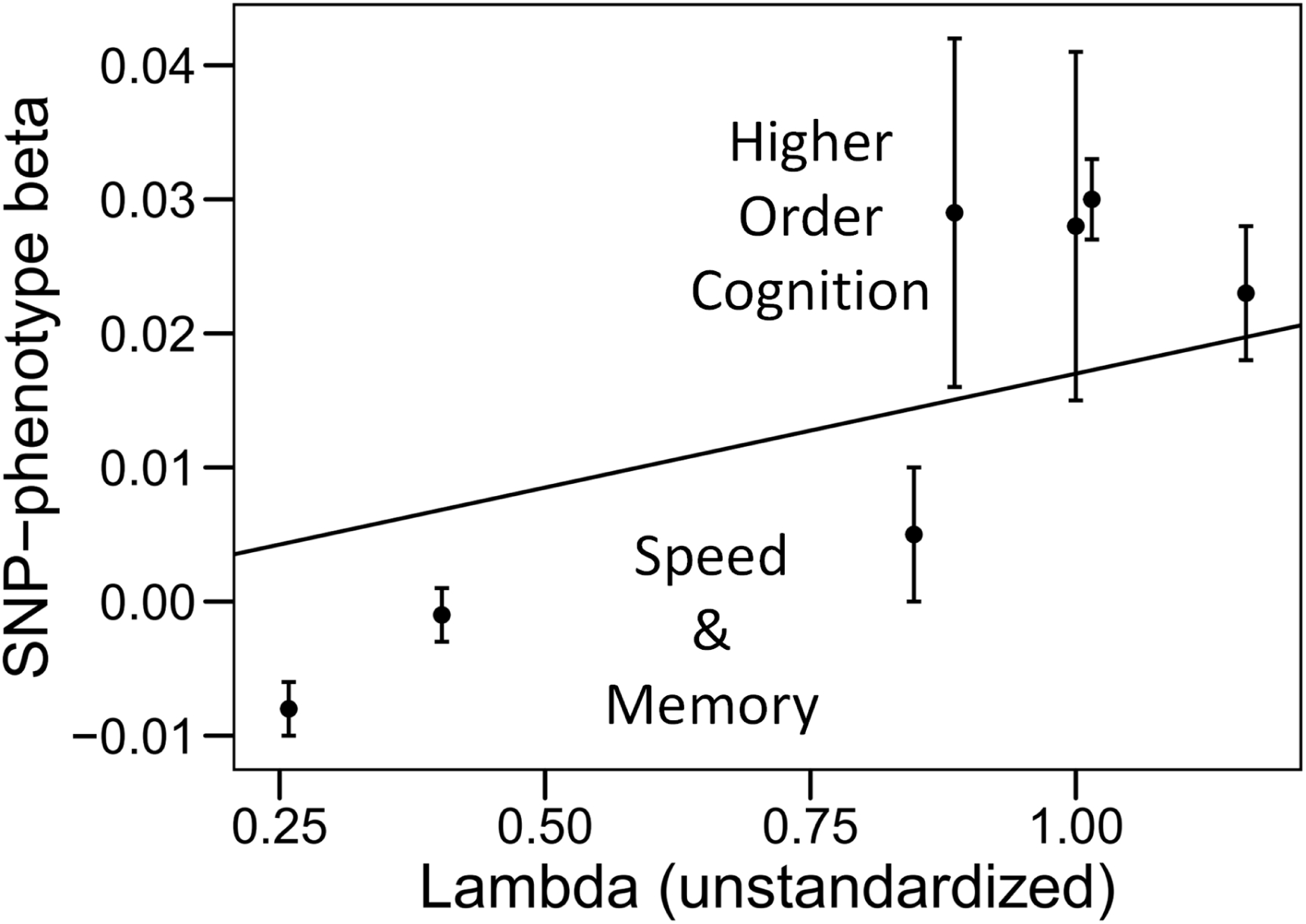
Scatter plot of SNP-phenotype regression coefficients against unstandardized genetic factor loadings for the associated phenotypes, for lead SNP **rs2352974** within a locus on Chromosome 3. Error bars represent standard errors of the SNP-phenotype betas. The solid black line represents the regression line based on all seven data points. In order to correspond directly with Equation S1, the regression line was estimated with its intercept fixed to zero, using weights equal to the inverse of the squared standard errors of the betas.

**Table S1.** SNP-based heritability (SNP h^2^; diagonal), LDSC-estimated genetic variance-covariance matrix (lower triangle) and genetic correlation matrix (upper triangle) across UK Biobank’s cognitive phenotypes.

|  | Matrix | Memory | RT | Symbol Digit | Trails-B | Tower | VNR |
| --- | --- | --- | --- | --- | --- | --- | --- |
| Matrix | 0.155 (0.040) | 0.473 (0.081) | 0.135 (0.071) | 0.544 (0.095) | 0.687 (0.094) | 0.633 (0.291) | 0.869 (0.069) |
| Memory | 0.037 (0.006) | 0.040 (0.002) | 0.218 (0.029) | 0.595 (0.046) | 0.618 (0.047) | 0.734 (0.095) | 0.429 (0.031) |
| RT | 0.014 (0.008) | 0.012 (0.002) | 0.074 (0.003) | 0.357 (0.035) | 0.312 (0.032) | 0.222 (0.072) | 0.169 (0.024) |
| Symbol Digit | 0.071 (0.012) | 0.040 (0.003) | 0.032 (0.003) | 0.110 (0.008) | 0.816 (0.056) | 0.665 (0.107) | 0.551 (0.034) |
| Trails-B | 0.104 (0.014) | 0.048 (0.004) | 0.033 (0.003) | 0.104 (0.007) | 0.149 (0.009) | 0.693 (0.104) | 0.743 (0.038) |
| Tower | 0.084 (0.029) | 0.050 (0.006) | 0.020 (0.007) | 0.074 (0.012) | 0.090 (0.014) | 0.114 (0.038) | 0.670 (0.080) |
| VNR | 0.157 (0.013) | 0.040 (0.003) | 0.021 (0.003) | 0.084 (0.005) | 0.132 (0.007) | 0.104 (0.012) | 0.212 (0.008) |

Matrix = Matrix Pattern Completion task; Memory = Memory – Pairs Matching Test; RT = Reaction Time; Symbol Digit = Symbol Digit Substitution Task; Trails-B = Trail Making Test – B; Tower = Tower Rearranging Task; VNR = Verbal Numerical Reasoning Test. The diagonal elements of the matrix contain SNP $h^2$ . The lower off-diagonal elements contain the genetic covariances across cognitive phenotypes, with standard errors in parentheses. The upper off-diagonal elements contain the genetic correlations across cognitive phenotypes, with standard errors in parentheses.

**Table S2.**
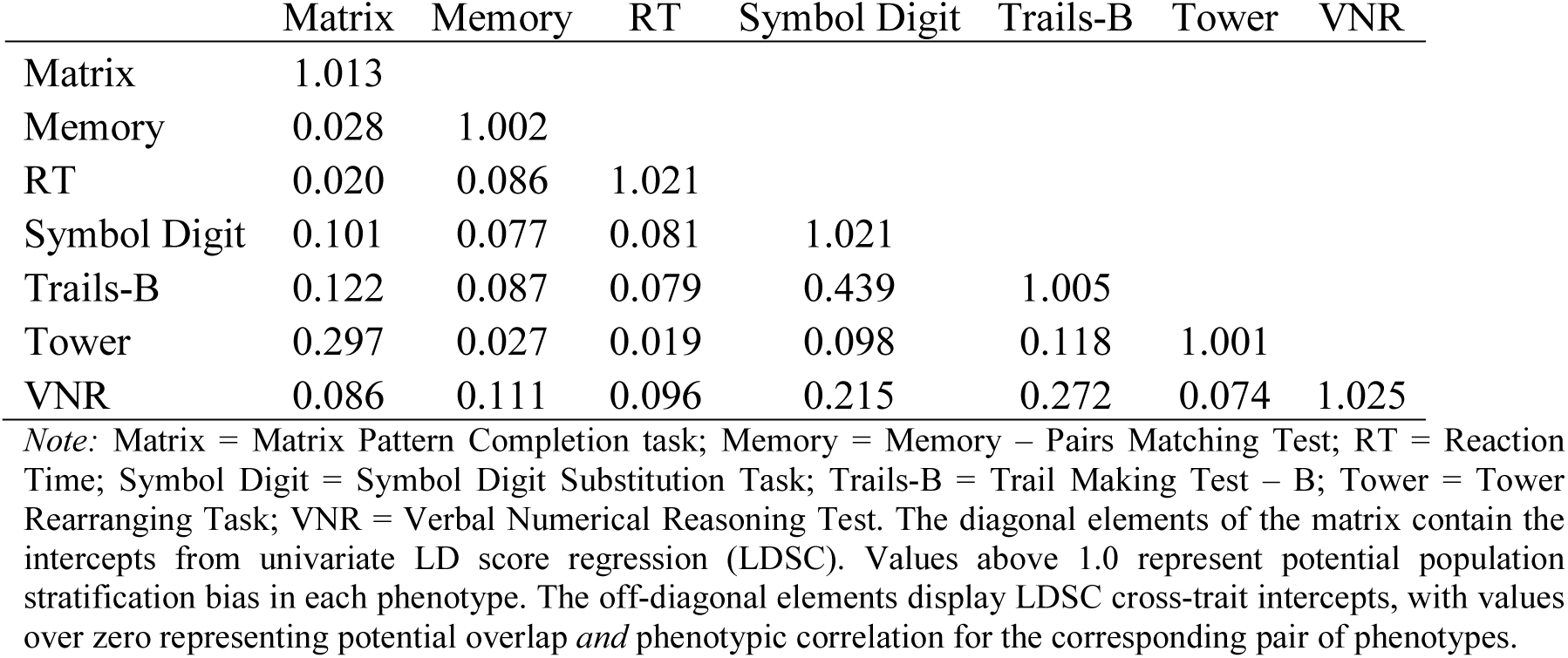
Intercepts and cross-trait intercepts from LDSC analysis of UK Biobank’s cognitive phenotypes.

**Table S3.** Phenotypic correlations across UK Biobank’s cognitive phenotypes.

|  | Matrix | Memory | RT | Symbol<br>Digit | Trails-B | Tower | VNR | *Note:*<br>Matr<br>ix =<br>Matr<br>ix<br>Patte<br>rn<br>Com<br>pleti<br>on<br>task;<br>Mem<br>ory =<br>Mem<br>ory –<br>Pairs |
| --- | --- | --- | --- | --- | --- | --- | --- | --- |
| Matrix | 1 |  |  |  |  |  |  |  |
| Memory | 0.178<br>(0.009) | 1 |  |  |  |  |  |  |
| RT | 0.184<br>(0.009) | 0.117<br>(0.002) | 1 |  |  |  |  |  |
| Symbol<br>Digit | 0.374<br>(0.011) | 0.199<br>(0.003) | 0.274<br>(0.003) | 1 |  |  |  |  |
| Trails-B | 0.412<br>(0.011) | 0.210<br>(0.004) | 0.272<br>(0.004) | 0.597<br>(0.003) | 1 |  |  |  |
| Tower | 0.353<br>(0.008) | 0.184<br>(0.009) | 0.192<br>(0.009) | 0.354<br>(0.012) | 0.394<br>(0.011) | 1 |  |  |
| VNR | 0.375<br>(0.014) | 0.140<br>(0.004) | 0.145<br>(0.005) | 0.347<br>(0.004) | 0.447<br>(0.004) | 0.305<br>(0.014) | 1 |  |

Matching Test; RT = Reaction Time; Symbol Digit = Symbol Digit Substitution Task; Trails-B = Trail Making Test – B; Tower = Tower Rearranging Task; VNR = Verbal Numerical Reasoning Test. The lower off-diagonal elements contain the phenotypic correlations across cognitive phenotypes, with standard errors in parentheses.

**Table S4.** Unstandardized and standardized common factor solutions for the genetic covariance structure of seven UK Biobank cognitive traits.

|  |  | Including Speeded Tests |  |  |  | Excluding Speeded Tests |  |  |  |
| --- | --- | --- | --- | --- | --- | --- | --- | --- | --- |
|  |  | Unstandardized | SE | Standardized | SE | Unstandardized | SE | Standardized | SE |
| Factor loadings |  |  |  |  |  |  |  |  |  |
|  | Matrix | 0.325 | 0.028 | 0.826 | 0.070 | 0.393 | 0.031 | 1.000 | 0.082 |
|  | Memory | 0.131 | 0.006 | 0.651 | 0.031 | 0.111 | 0.008 | 0.555 | 0.040 |
|  | RT | 0.084 | 0.007 | 0.308 | 0.026 | - | - | - | - |
|  | Symbol Digit | 0.275 | 0.011 | 0.831 | 0.034 | - | - | - | - |
|  | Trails-B | 0.377 | 0.014 | 0.976 | 0.035 | - | - | - | - |
|  | Tower | 0.288 | 0.027 | 0.853 | 0.080 | 0.310 | 0.030 | 0.921 | 0.089 |
|  | VNR | 0.330 | 0.011 | 0.717 | 0.024 | 0.364 | 0.022 | 0.793 | 0.048 |
| Residual variances |  |  |  |  |  |  |  |  |  |
|  | Matrix | 0.049 | 0.038 | 0.317 | 0.243 | 0.000 | 0.039 | 0.000 | 0.251 |
|  | Memory | 0.023 | 0.002 | 0.576 | 0.050 | 0.028 | 0.002 | 0.692 | 0.061 |
|  | RT | 0.067 | 0.003 | 0.905 | 0.043 | - | - | - | - |
|  | Symbol Digit | 0.034 | 0.006 | 0.309 | 0.050 | - | - | - | - |
|  | Trails-B | 0.007 | 0.008 | 0.048 | 0.051 | - | - | - | - |
|  | Tower | 0.031 | 0.034 | 0.272 | 0.299 | 0.017 | 0.035 | 0.151 | 0.310 |
|  | VNR | 0.103 | 0.007 | 0.487 | 0.033 | 0.079 | 0.016 | 0.371 | 0.076 |

*Note:* Matrix = Matrix Pattern Completion task; Memory = Memory – Pairs Matching Test; RT = Reaction Time; Symbol Digit = Symbol Digit Substitution Task; Trails-B = Trail Making Test – B; Tower = Tower Rearranging Task; VNR = Verbal Numerical Reasoning Test. Standardized factor loadings indicate the lineal relationship between the genetic *g* factor and each of the cognitive phenotypes, ranging from -1 to 1, with 0 representing no relationship. SE = Standard Error. $R^2$ = percentage of genetic variance of each phenotype accounted for the genetic *g* factor. Residual variances reflect genetic variation unique to each cognitive phenotype. Note: In the model that excluded speeded tests, there was a Heywood case for the Matrix loading. Its residual variance was subsequently constrained to be greater than 0.

**Table S5.**
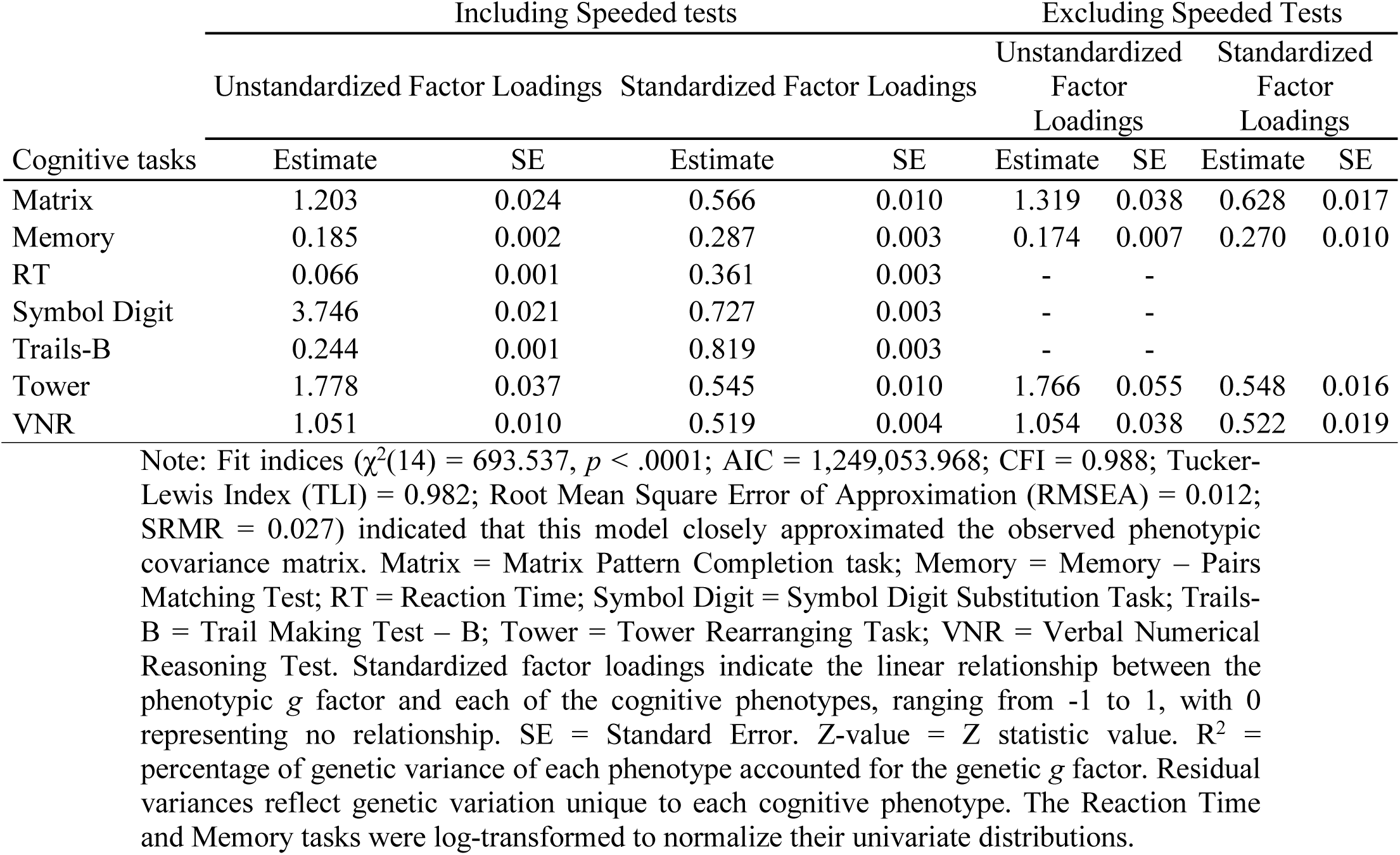
Phenotypic *g* factor CFA estimates for UK Biobank’s cognitive phenotypes.

| Cognitive tasks | Including Speeded tests |  |  |  | Excluding Speeded Tests |  |  |  |
| --- | --- | --- | --- | --- | --- | --- | --- | --- |
|  | Unstandardized Factor Loadings |  | Standardized Factor Loadings |  | Unstandardized Factor Loadings |  | Standardized Factor Loadings |  |
|  | Estimate | SE | Estimate | SE | Estimate | SE | Estimate | SE |
| Matrix | 1.203 | 0.024 | 0.566 | 0.010 | 1.319 | 0.038 | 0.628 | 0.017 |
| Memory | 0.185 | 0.002 | 0.287 | 0.003 | 0.174 | 0.007 | 0.270 | 0.010 |
| RT | 0.066 | 0.001 | 0.361 | 0.003 | - | - |  |  |
| Symbol Digit | 3.746 | 0.021 | 0.727 | 0.003 | - | - |  |  |
| Trails-B | 0.244 | 0.001 | 0.819 | 0.003 | - | - |  |  |
| Tower | 1.778 | 0.037 | 0.545 | 0.010 | 1.766 | 0.055 | 0.548 | 0.016 |
| VNR | 1.051 | 0.010 | 0.519 | 0.004 | 1.054 | 0.038 | 0.522 | 0.019 |

Note: Fit indices ( $\chi^2(14) = 693.537$ , $p < .0001$ ; AIC = 1,249,053.968; CFI = 0.988; Tucker-Lewis Index (TLI) = 0.982; Root Mean Square Error of Approximation (RMSEA) = 0.012; SRMR = 0.027) indicated that this model closely approximated the observed phenotypic covariance matrix. Matrix = Matrix Pattern Completion task; Memory = Memory – Pairs Matching Test; RT = Reaction Time; Symbol Digit = Symbol Digit Substitution Task; Trails-B = Trail Making Test – B; Tower = Tower Rearranging Task; VNR = Verbal Numerical Reasoning Test. Standardized factor loadings indicate the linear relationship between the phenotypic *g* factor and each of the cognitive phenotypes, ranging from -1 to 1, with 0 representing no relationship. SE = Standard Error. Z-value = Z statistic value. $R^2$ = percentage of genetic variance of each phenotype accounted for the genetic *g* factor. Residual variances reflect genetic variation unique to each cognitive phenotype. The Reaction Time and Memory tasks were log-transformed to normalize their univariate distributions.

**Table S6.** Genetic correlations of genetic *g* derived from the UK Biobank and genetic results from other major intelligence studies with educational attainment, neural phenotypes, and longevity.

|  | Genetic *g*<br>(speeded tests<br>included):<br>Present Study |  | Genetic *g*<br>(speeded tests<br>excluded):<br>Present Study |  | General Cognitive<br>Function:<br>Davies et al. (2018) |  | Intelligence:<br>Savage et al. (2019) |  | Intelligence:<br>Hill et al. (2019) |  |
| --- | --- | --- | --- | --- | --- | --- | --- | --- | --- | --- |
|  | *r* | SE | *r* | SE | *r* | SE | *r* | SE | *r* | SE |
| Educational Attainment:<br>(Lee et al., 2018) | 0.475 | 0.023 | 0.596 | 0.021 | 0.694 | 0.027 | 0.730 | 0.034 | 0.847 | 0.027 |
| Alzheimer's Disease:<br>Lambert et al. (2013) | -0.341 | 0.054 | -0.329 | 0.064 | -0.326 | 0.049 | -0.290 | 0.068 | -0.327 | 0.046 |
| Autism Spectrum Disorder:<br>Brainstorm Consortium<br>(2018) | 0.095 | 0.043 | 0.173 | 0.043 | 0.203 | 0.036 | 0.234 | 0.038 | 0.205 | 0.034 |
| ADHD:<br>Brainstorm Consortium<br>(2018) | -0.233 | 0.038 | -0.323 | 0.040 | -0.376 | 0.035 | -0.379 | 0.042 | -0.462 | 0.034 |
| Schizophrenia:<br>Brainstorm Consortium<br>(2018) | -0.374 | 0.029 | -0.340 | 0.031 | -0.264 | 0.021 | -0.233 | 0.029 | -0.166 | 0.020 |
| Total Brain Volume:<br>Zhao et al. (2019) | 0.195 | 0.041 | 0.253 | 0.046 | 0.229 | 0.040 | 0.229 | 0.051 | 0.250 | 0.040 |
| Longevity:<br>Timmers et al. (2019) | 0.255 | 0.030 | 0.290 | 0.031 | 0.320 | 0.027 | 0.284 | 0.036 | 0.377 | 0.026 |

*Note:* all correlations are statistically significant with $p < .001$ , except $r$ Genetic *g* with ASD ( $p = .029$ ). ADHD = Attention Deficit Hyperactivity Disorder. $r$ = Pearson correlation coefficient. SE = Standard Error.

